# Structural basis for Cas9 off-target activity

**DOI:** 10.1101/2021.11.18.469088

**Authors:** Martin Pacesa, Chun-Han Lin, Antoine Cléry, Aakash Saha, Pablo R. Arantes, Katja Bargsten, Matthew J. Irby, Frédéric H.T. Allain, Giulia Palermo, Peter Cameron, Paul D. Donohoue, Martin Jinek

## Abstract

The target DNA specificity of the CRISPR-associated genome editor nuclease Cas9 is determined by complementarity to a 20-nucleotide segment in its guide RNA. However, Cas9 can bind and cleave partially complementary off-target sequences, which raises safety concerns for its use in clinical applications. Here we report crystallographic structures of Cas9 bound to *bona fide* off-target substrates, revealing that off-target binding is enabled by a range of non-canonical base pairing interactions within the guide–off-target heteroduplex. Off-target sites containing single-nucleotide deletions relative to the guide RNA are accommodated by base skipping or multiple non-canonical base pairs rather than RNA bulge formation. Additionally, PAM-distal mismatches result in duplex unpairing and induce a conformational change of the Cas9 REC lobe that perturbs its conformational activation. Together, these insights provide a structural rationale for the off-target activity of Cas9 and contribute to the improved rational design of guide RNAs and off-target prediction algorithms.

## INTRODUCTION

Cas9, the effector nuclease of prokaryotic Type II CRISPR adaptive immune systems (Makarova et al., 2020), cleaves double-stranded DNA substrates complementary to a guide CRISPR RNA (crRNA) (Jinek et al., 2012). By changing the sequence of the guide RNA, the target DNA specificity of the CRISPR-Cas9 system is readily programmable (Jinek et al., 2012), a feature that has been widely exploited for genome engineering applications (Anzalone et al., 2020). Cas9 functions in conjunction with a trans-activating crRNA (tracrRNA), which is required both for crRNA loading and subsequent DNA binding and cleavage (Deltcheva et al., 2011; Jinek et al., 2012). Target DNA binding and cleavage is further dependent on the presence of a protospacer-adjacent motif (PAM) flanking the target sequence (Anders et al., 2014; Jinek et al., 2012). Due to its high activity and 5’-NGG-3’ PAM specificity, *Streptococcus pyogenes* Cas9 (SpCas9) remains the most widely used CRISPR-Cas nuclease for gene editing applications. However, despite a high intrinsic accuracy in generating targeted DNA breaks, SpCas9 can nevertheless cleave genomic DNA sequences with imperfect complementarity to the guide RNA, resulting in off-target editing (Cameron et al., 2017; Hsu et al., 2013; Pattanayak et al., 2013; Tsai et al., 2015). The off-target activity of SpCas9, as well as other Cas9 enzymes, thus presents a safety concern for their therapeutic applications.

Off-target sites typically contain one or several nucleobase mismatches relative to the guide RNA (Cameron et al., 2017; Tsai et al., 2017; Tsai et al., 2015). Recent studies have established that the type of mismatch, its positioning, and the total number of mismatches are important determinants of off-target DNA binding and cleavage (Boyle et al., 2017; Boyle et al., 2021; Doench et al., 2016; Jones et al., 2021; Zhang et al., 2020). PAM-proximal mismatches within the seed region of the guide RNA (gRNA)-target strand (TS) DNA heteroduplex typically have a dramatic impact on substrate DNA binding and R-loop formation (Boyle et al., 2021; Ivanov et al., 2020; Singh et al., 2016; Zhang et al., 2020). In contrast, PAM-distal mismatches are compatible with stable binding; however, their presence often results in the formation of a catalytically incompetent complex (Boyle et al., 2021; Dagdas et al., 2017; Ivanov et al., 2020; Jones et al., 2021; Sternberg et al., 2015; Yang et al., 2018; Zhang et al., 2020). In addition, Cas9 has been shown to cleave off-target substrates containing insertions or deletions relative to the guide RNA sequence, which have been proposed to be recognized through the formation of nucleotide “bulges” in the gRNA-TS DNA heteroduplex (Boyle et al., 2021; Cameron et al., 2017; Doench et al., 2016; Jones et al., 2021; Lin et al., 2014; Tsai et al., 2015).

Numerous computational tools have been developed to predict possible genomic off-target sites based on sequence similarity (Bae et al., 2014; Stemmer et al., 2015). However, the majority of actual off-target cleavage events remain unpredicted (Cameron et al., 2017; Tsai et al., 2015). Furthermore, although Cas9 is able to bind genomic sites harbouring as few as five complementary nucleotides, only a relatively small number of off-target sites are actually cleaved and result in detectable off-target editing in cells (Kuscu et al., 2014; O’Geen et al., 2015; Wu et al., 2014). Several structures of target-bound Cas9 complexes have been determined to date (Anders et al., 2014; Jiang et al., 2016; Nishimasu et al., 2014; Zhu et al., 2019) that have shed light on the mechanism of on-target binding and cleavage. However, the same processes for off-target sites remain poorly understood.

To elucidate the mechanism of mismatch tolerance of Cas9, we determined crystal structures of a comprehensive set of *bona fide* off-target–bound complexes. These structures reveal that the formation of non-canonical base pairs and preservation of heteroduplex shape underpin the off-target tolerance of Cas9. We also observe that multiple consecutive mismatches can be accommodated by base skipping of a guide RNA nucleotide, as opposed to nucleotide bulging. Finally, the structure of an off-target complex containing three PAM-distal mismatches exhibits REC2/3 domain rearrangements, which likely perturbs conformational activation of Cas9 and thus modulates cleavage efficiency. Taken together, our structural data reveal the diversity of mechanisms enabling off-target recognition and lay the foundation for improved off-target prediction and engineering optimized CRISPR-Cas9 complex designs for gene editing.

## RESULTS

### *In vitro* profiling reveals diversity of Cas9 off-targets

Multiple studies have investigated the off-target activity of Cas9, suggesting context-dependent tolerance of nucleobase mismatches between the guide RNA and off-target DNA sequences (Boyle et al., 2021; Cameron et al., 2017; Lazzarotto et al., 2020; Tsai et al., 2015; Zhang et al., 2020). To investigate the effect of mismatches on Cas9 binding and cleavage, we performed the SITE-Seq^®^ assay (Cameron et al., 2017) to define the off-target landscapes of 12 well-studied guide RNAs to select suitable off-targets for further evaluation (**Table S1, Table S2**) (**Figure S1A, Table 1**). The SITE-Seq assay analysis revealed a total of 3,848 detectable off-target sites at the highest Cas9 ribonucleoprotein (RNP) concentration, with a median of 5 mismatches per off-target site (**Figure S1C**). The detected mismatches covered all possible base mismatch combinations and were distributed throughout the length of the gRNA-TS DNA heteroduplex (**Figure S1D-E**).

**Table 1.**
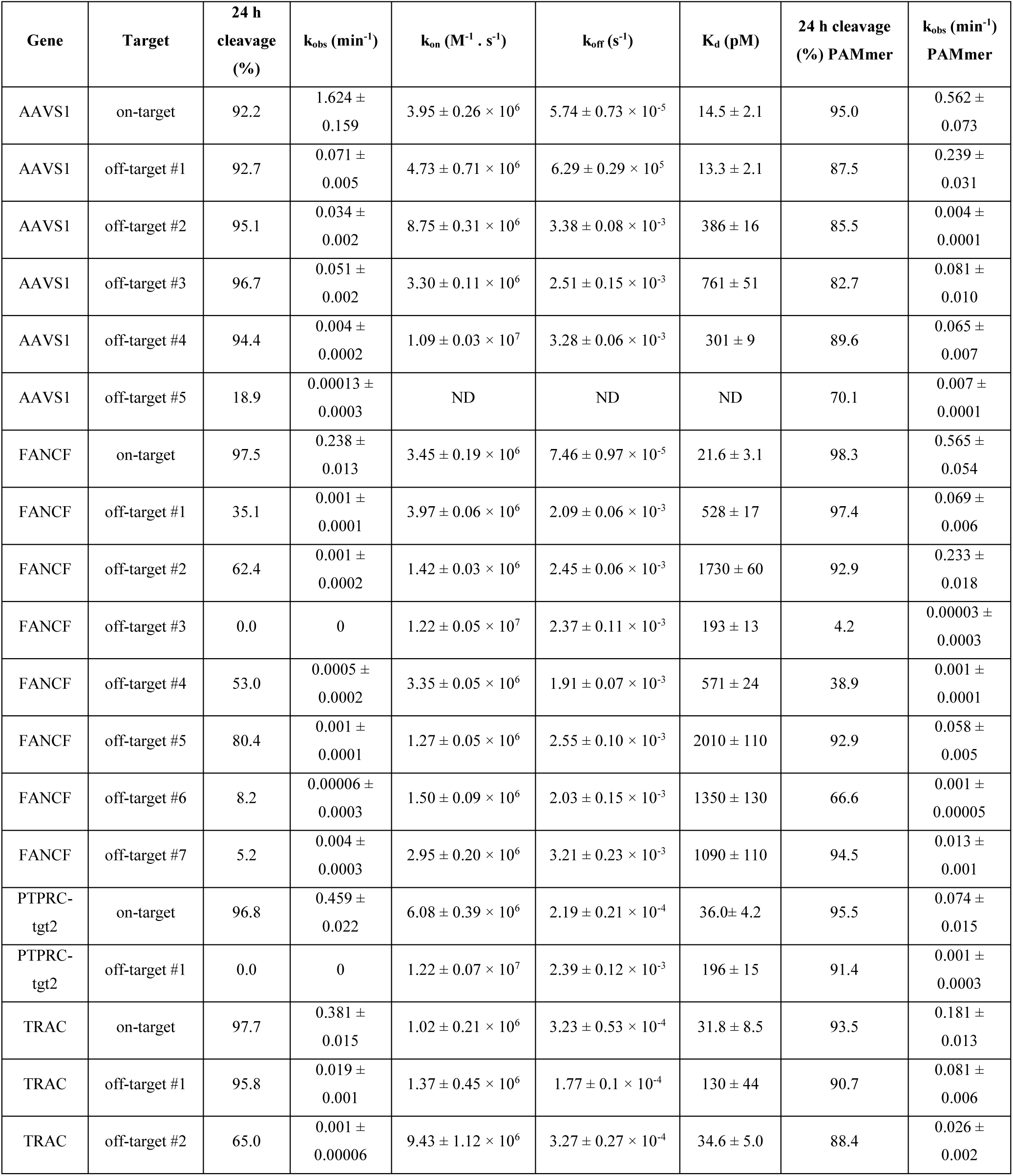
Kinetic and thermodynamic analysis of off-target substrate binding and cleavage. Kinetic and thermodynamic parameters of off-target substrates. The cleavage rate constants (k_obs_) were derived from single-exponential function fitting of measured cleavage rates of the target-strand labelled DNA substrates. The binding and dissociation rate constants (k_on_ and k_off_) and the equilibrium dissociation constant (K_d_) were determined using a DNA nanolever binding (switchSENSE) assay. Intervals indicate standard error of mean.

To probe the thermodynamics of on- and off-target substrate DNA binding and the kinetics of DNA cleavage, we focused on a subset of four guide RNAs (*AAVS1*, *FANCF*, *PTPRC-tgt2,* and *TRAC*) and a total of 15 *bona fide* off-target sites detectable *in vivo* (Cameron et al., 2017; Donohoue et al., 2021; Tsai et al., 2017; Tsai et al., 2015) (**Figure 1A**) that covered all 12 possible base mispair types. To benchmark the editing efficiency of the selected off-target substrates *in vivo*, we transfected human primary T cells with recombinant Cas9 RNPs at multiple concentrations and quantified indel frequencies. We observed efficient cellular editing at the on-target site for each guide RNA at all RNP concentrations, and detected editing at 7 of the 15 off-target sites in our set, including the deletion-containing FANCF off-target #3 (**Figure S3, Table S3**), which contained a mismatch-reverting single-nucleotide polymorphism (SNP) relative to the consensus human genome build.

**Figure 1.**
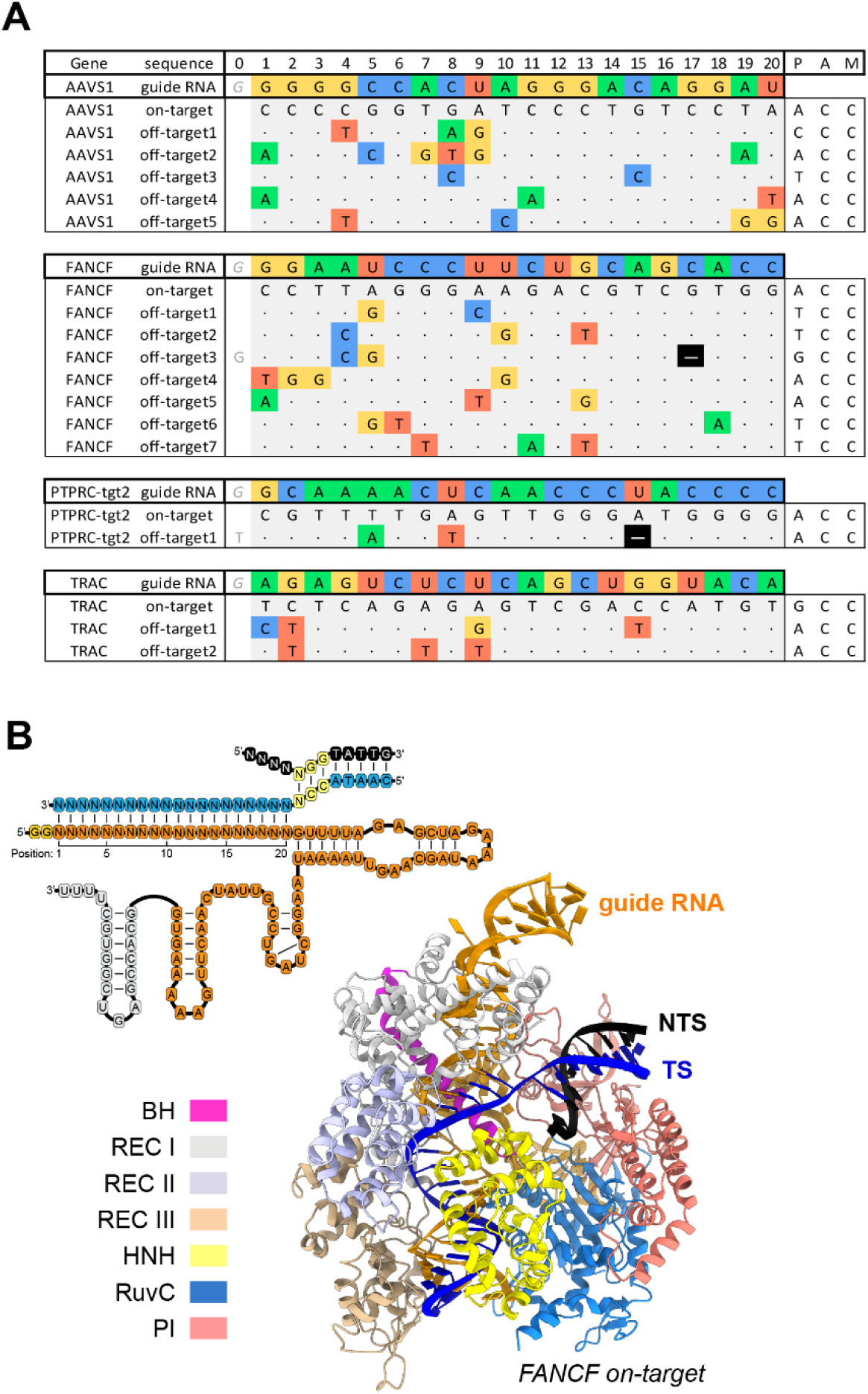
Biochemical and structural analysis of Cas9 off-targets. (**A**) Guide RNA and (off-)target DNA sequences selected for biochemical and structural analysis. Matching bases in off-targets are denoted by a dot; nucleotide mismatches and deletions (–) are highlighted. (**B**) Top: Schematic representation of the guide RNA (orange), TS (blue), and NTS (black) sequences used for crystallisation. The PAM sequence in the DNA is highlighted in yellow. Bottom: Structure of the Cas9 *FANCF* on-target complex. Individual Cas9 domains are coloured according to the legend; substrate DNA target strand (TS) is coloured blue, non-target strand (NTS) black, and the guide RNA orange.

*In vitro* nuclease activity assays revealed that all selected off-target sequences were cleaved slower than the corresponding on-target substrates, with 20–500-fold reductions in the calculated rate constants (**Table 1, Figure S2A**). To distinguish whether the cleavage defects were due to slow R-loop formation or perturbations in downstream steps, we also quantified cleavage kinetics using partially single-stranded PAMmer DNA substrates (Anders et al., 2014; O’Connell et al., 2014). These experiments revealed that the slower cleavage kinetics of most off-target substrates was due to perturbed R-loop formation (**Table 1, Figure S2B**). However, for some off-targets, notably *AAVS1* off-targets #2 and #5, *FANCF* off-targets #3, #4, and #6, and *PTPRC-tgt2* off-target #1, the rate of PAMmer substrate cleavage was more than 100-fold slower as compared to their respective on-target sequences (**Table 1**, **Figure S2B**), indicating perturbations in the conformational activation checkpoint downstream of guide-target hybridization or inhibition of cleavage by direct steric hindrance of the Cas9 HNH domain (Chen et al., 2017; Dagdas et al., 2017).

Complementary quantification of substrate DNA binding using DNA nanolever (switchSENSE) methodology revealed perturbations in the binding affinities of most off-targets (**Table 1**). Notably, the reductions in binding affinities were almost entirely due to increased dissociation rates (k_off_), while on-binding rates (k_on_) were largely unperturbed (**Table 1**), indicating that most of the tested off-targets promote DNA dissociation, likely due to R-loop collapse. However, there was little correlation between the observed reductions in cleavage rates and binding constants (**Figure S2C**), confirming that the molecular basis for off-target discrimination by Cas9 is not based on substrate binding alone, in agreement with prior studies (Boyle et al., 2021; Chen et al., 2017; Dagdas et al., 2017; Jones et al., 2021; Yang et al., 2018; Zhang et al., 2020). The dissociation rate (k_off_) correlated significantly only with the number of mismatches located in the seed region (R^2^=0.46, p=0.001) (**Figure S2C**), suggesting that seed mismatches promote R-loop collapse and non-target strand (NTS) rehybridization (Boyle et al., 2017; Gong et al., 2018; Singh et al., 2016; Sternberg et al., 2014).

Finally, we compared the measured cleavage rate constants (k_obs_) with predicted data from a leading biophysical model of Cas9 off-target cleavage that accounts for mismatch number and position but utilises position-independent weights for mismatch type (Jones et al., 2021). Despite good overall correlation between the model and our data (R^2^=0.46, p=0.004) (**Figure S2C**), there were several prominent outliers (*AAVS1* off-target #2 and off-target #3, and *FANCF* off-target #4), suggesting that accurate modelling of off-target interactions requires accounting for position-specific effects of individual mismatch types (**Figure S2D**).

### Crystallographic analysis of off-target interactions

To obtain insights into the structural basis of off-target recognition and mismatch tolerance, we employed a previously described approach (Anders et al., 2014) to co-crystallize Cas9 with sgRNA guides and partially duplexed off-target DNA substrates (**Figure 1C**). Focusing on our set of *AAVS1*, *FANCF*, *PTPRC-tgt2* and *TRAC* off-targets (**Figure 1A**), covering all 12 possible mismatch types, we determined a total of 15 off-target complex structures at resolutions of 2.25–3.30 Å (**Figure 1C, Table S4**). Overall, the off-target complex structures have very similar conformations, with the Cas9 polypeptide backbone superimposing with a mean root mean square deviation of 0.41Å over 1330 Cα atoms (as referenced to the *FANCF* on-target complex structure, excluding *FANCF* off-target #4, as discussed below). Of note, the *AAVS1* on-target complex structure reveals substantial repositioning of the REC2 domain, where it undergoes a 12° rotation (relative to *FANCF* and *TRAC* on-target complexes) (**Figure S4A**), with concomitant shortening of the α-helix comprising residues 301-305 and restructuring of the loop comprising residues 175-179 (**Figure S4B**), enabled by the absence of crystal contacts involving the REC2 and REC3 domains.

However, the structures display considerable local variation of the gRNA-TS DNA heteroduplex conformation (**Table S5**). Base pairing and base stacking are mostly preserved throughout the heteroduplexes (**Table S6**), with the exception of positions 1–3 within the PAM-distal end of the guide-TS duplex, where the presence of mismatches results in duplex unpairing (**Figure S5**). Interestingly, such unpairing results in the base stacking of the terminal residue of the sgRNA on top of the unpaired DNA base, as observed in a recent cryo-EM structure of a PAM-distal mismatch off-target complex (Bravo et al., 2022). Despite the observed conformational variation, the off-target structures preserve almost all intermolecular contacts between the Cas9 protein and the bound nucleic acids (**Figure S6**), further underscoring the structural plasticity of Cas9 in accommodating mismatch-induced heteroduplex distortions.

### Non-canonical base-pairing interactions facilitate off-target recognition

A substantial fraction of the base pair mismatches observed in the determined off-target complex structures (34 out of 49) is accommodated by non-canonical base pairing interactions that preserve at least one hydrogen bond between the mispaired bases. The most common off-target mismatches, both in our data set (**Table S1, Table S2**) and as reported by other studies (Boyle et al., 2017; Doench et al., 2016; Hsu et al., 2013; Jones et al., 2021; Pattanayak et al., 2013; Zhang et al., 2020), are rG-dT (**Figure S7**) and rU-dG (**Figure 2A, Figure S8**), which have the potential to form wobble base pairs (Kimsey et al., 2015). Indeed, all rG-dT mismatches in the determined structures are accommodated by wobble base pairing. Observed at duplex positions 4, 13 and 15, the dT base undergoes a ~1 Å shear displacement into the major groove of the gRNA–TS DNA heteroduplex to form the wobble base pair (**Figure S7A-E**), whereas at duplex position 2, wobble base pairing is enabled by a minor groove displacement of the rG base (**Figure S7F-G**). In contrast, rU-dG base pairs in the determined structures exhibit considerable structural variation. At duplex position 10, the rU base is able to undergo the major groove displacement required for wobble base-pairing (**Figure 2A, Figure S8A**). In contrast, at duplex position 5, the backbone of the RNA strand makes extensive contacts with Cas9 (**Figure S8B-D, Figure S6**) and as a result, the rU-dG base pairs are instead accommodated by compensatory shifts of the dG base to maintain hydrogen-bonding interactions (**Figure S8B-D**). At duplex position 9 in the *TRAC* off-target #1 complex, the rU-dG mismatch is accommodated by wobble base-pairing enabled by a minor groove displacement of the dG base (**Figure S8E**). At the same duplex position in the *AAVS1* off-target #1 and #2 complexes, however, this mismatch occurs next to rC-dA and rC-dT mismatches, respectively, and adopts the sterically prohibited Watson-Crick geometry (**Figure S8F-G**), implying a tautomeric shift or base deprotonation to accommodate this otherwise unfavorable base pairing mode (**Figure S8H**). Collectively, these observations suggest that the ability of rU-dG (and likely rG-dT) mismatches to form wobble base pairing interactions is determined not only by backbone constraints at the specific position within the gRNA-TS DNA heteroduplex (**Figure S6**), but also by local sequence context and/or the presence of neighboring mismatches (**Figure S8C-G**).

**Figure 2.**
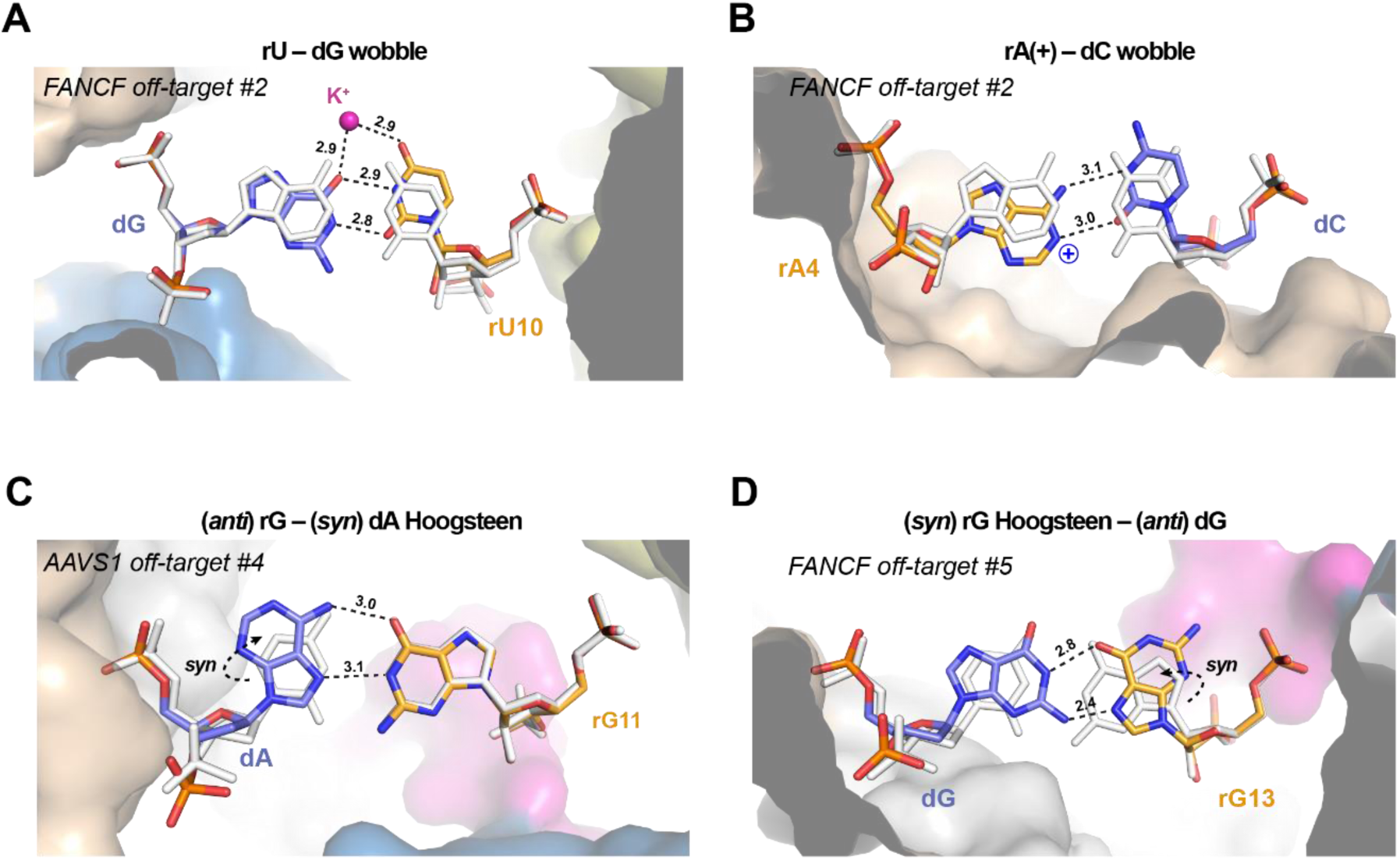
Cas9 off-target binding is enabled by non-canonical base pairing. Close-up views of (**A**) rU-dG wobble base pair at duplex position 10 in *FANCF* off-target #2 complex, (**B**) rA-dC wobble base pair at position 4 in *FANCF* off-target #2 complex, (**C**) rG-dA Hoogsteen base pair at duplex position 11 in *AAVS1* off-target #4 complex and (**D**) rG-dG Hoogsteen base pair at duplex position 13 in *FANCF* off-target #5 complex. Hydrogen bonding interactions are indicated with dashed lines. Numbers indicate interatomic distances in Å. Corresponding on-target Watson-Crick base pairs are shown in white. Dashed arrows indicate *anti*-*syn* isomerization of the dA and rG bases to enable Hoogsteen-edge base pairing. A bound monovalent ion, modelled as K^+^, is depicted as a purple sphere.

rA-dC or rC-dA mismatches can also form wobble-like base pairs when the adenine base is protonated at the N1 position (Garg and Heinemann, 2018; Wang et al., 2011). In the rA-dC mispair found at duplex position 4, the dC base undergoes a wobble displacement compatible with the formation of two hydrogen bonds with the adenine base, indicative of adenine protonation (**Figure 2B**, **Figure S9A**). At other duplex positions in our data set, the rA-dC or rC-dA mispairs are instead accommodated by slight displacements of the adenine base within the base pair plane resulting in the formation of a single hydrogen bond in each case (**Figure S9B-D**).

Accommodating purine-purine mismatches by Watson-Crick-like interactions would normally require severe distortion of the guide–off-target duplex to increase its width by more than 2 Å (Leontis et al., 2002). At positions where the duplex width is constrained by Cas9 interactions (**Figure S6**), rG-dA and rA-dG mispairs are accommodated by *anti*-to-*syn* isomerization of the adenine base to form two hydrogen-bonding interactions via its Hoogsteen base edge. This is observed at duplex position 11 in the *AAVS1* off-target #4 complex (rG-dA mispair) (**Figure 2C**) and at position 7 in the *AAVS1* off-target #2 complex (rA-dG mispair) (**Figure S9E**). Similarly, the rG-dG mispair at duplex position 13 in the *FANCF* off-target #5 complex is accommodated by Hoogsteen base-pairing as a result by *anti*-to-*syn* isomerization of the guide RNA base (**Figure 2D**). Overall, the observed Hoogsteen base pairing interactions are near-isosteric with Watson-Crick base pairs and maintain duplex width without excessive backbone distortion (**Table S6**).

### Duplex backbone rearrangements accommodate otherwise non-productive mismatches

Whereas wobble (G-U/T or A-C) and Hoogsteen (A-G or G-G) base pairs are generally compatible with the canonical A-form geometry of an RNA-DNA duplex, other nucleotide mismatches only form non-isosteric base pairs that require considerable distortion of the (deoxy)ribose-phosphate backbone. The formation of pyrimidine-pyrimidine base pairs is expected to occur by a substantial reduction in duplex width (Leontis et al., 2002). This is observed at the rU-dC mismatch at duplex position 9 in the *FANCF* off-target #1 complex (**Figure 3A**). Here, the guide RNA backbone is able to shift towards the target DNA strand, resulting in a reduction of the C1’–C1’ distance to 8.65 Å as compared to 10.0 Å in the *FANCF* on-target complex. This facilitates the formation of two hydrogen bonding interactions within the rU-dC base pair, which is further enabled by a substantial increase in base propeller twist (**Figure 3A**). In contrast, rC-dT mismatches remain unpaired at duplex positions 6 and 7 in (**Figure S10A-B**), or form only a single hydrogen bond at position 8 (**Figure S10C**), likely due to backbone steric constraints at these positions imposed by Cas9 interactions (**Figure S6**). Of note, the *FANCF* off-target #7 rC-dT mismatch is bridged by hydrogen bonding interactions with the side chain of Arg895 inserted into the minor groove of the heteroduplex (**Figure S10B**); however, the interaction is not essential for the tolerance of rC-dT mismatches at this position (**Figure S11, Table S7**). Backbone steric constraints also likely influence the formation of rU-dT base pairs (**Figure S6**). At duplex position 7, the mismatch remains unpaired (**Figure S10D**), whereas productive pairing is seen at duplex positions 8 and 9, facilitated by distortions of the guide RNA and TS backbone, respectively (**Figure 3B, Figure S10E-F**).

**Figure 3.**
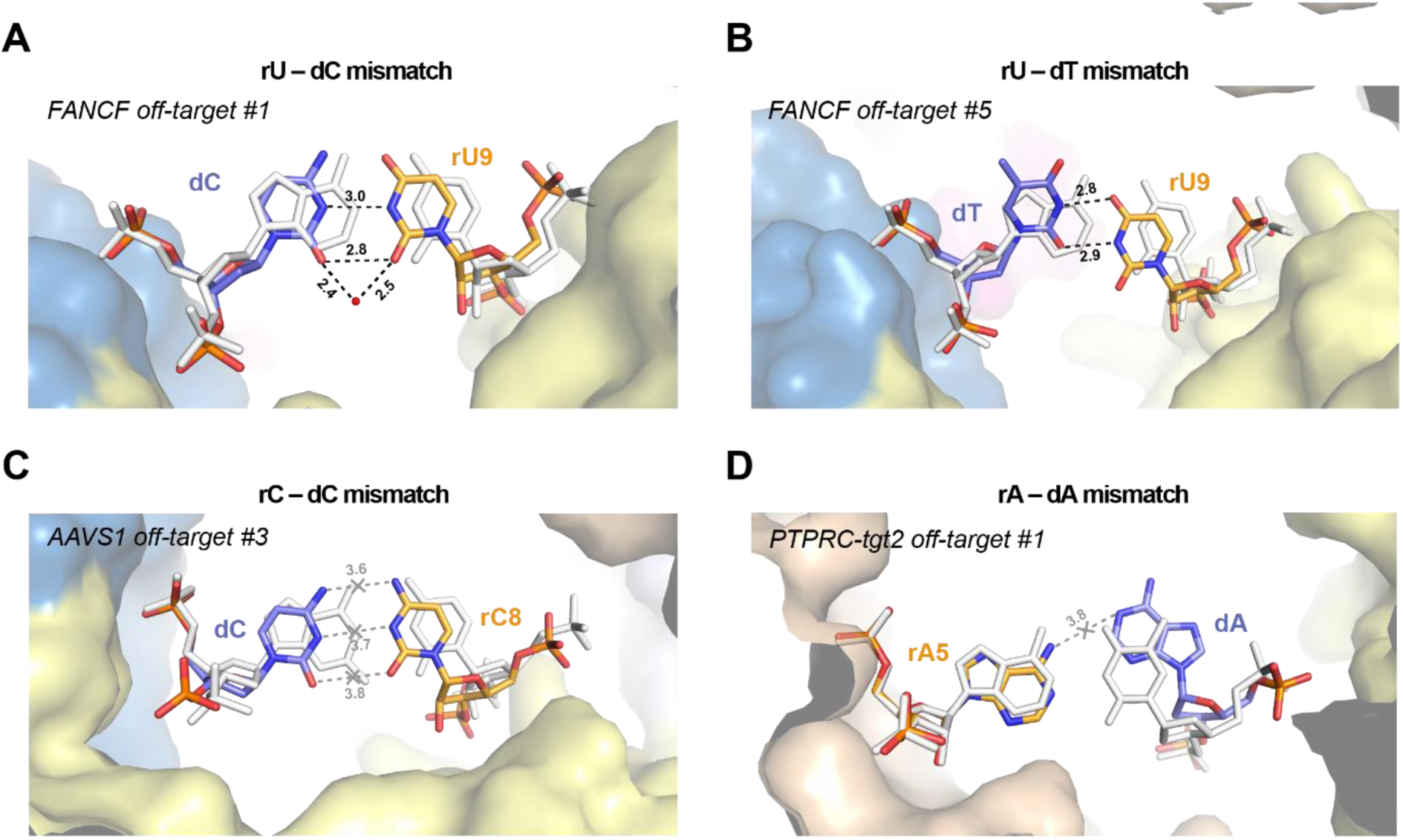
Duplex backbone distortions facilitate formation of non-canonical base pairs. (**A**) Close-up view of the rU-dC base pair at duplex position 9 in *FANCF* off-target #1 complex, facilitated by lateral displacement of the guide RNA backbone. (**B**) Zoomed-in view of the rU-dT base pair at position 9 in *FANCF* off-target #5 complex. (**C**) Zoomed-in view of the rC-dC mismatch at duplex position 8 in *AAVS1* off-target #3 complex. The distances between the cytosine bases indicate lack of hydrogen bonding. (**D**) Zoomed-in view of the rA-dA mismatch at duplex position 5 in *PTPRC-tgt2* off-target #1 complex. Hydrogen bonding interactions are indicated with dashed lines. Corresponding on-target Watson-Crick base pairs are shown in white (for *PTPRC-tgt2*, the *FANCF* on-target structure was used and bases were mutated *in silico*). Numbers indicate interatomic distances in Å. Bound water molecule is depicted as red sphere.

rC-dC mismatches only form productive hydrogen-bonding interactions if bridged by a water molecule or when one of the cytosine bases is protonated (Leontis et al., 2002). Only the former is observed in the determined structures, at duplex position 5 in the *AAVS1* off-target #2 complex (**Figure S10G**). In contrast, at duplex position 8 and 15, the bases remain unpaired while maintaining intra-strand base stacking interactions (**Figure 3C, Figure S10H**). Similarly, rA-dA mismatches are unable to form productive hydrogen-bonding interactions within the constraints of an A-form duplex (Leontis et al., 2002). Accordingly, the rA-dA mismatch at duplex position 5, where duplex width is constrained by Cas9 (**Figure S6**), is accommodated by extrusion of the dA nucleobase out of the base stack into the major groove of the duplex (**Figure 3D**) which is enabled by local distortion of the TS backbone (**Figure S12A**). Analysis of our SITE-Seq assay data set revealed that off-target rA-dA mismatches occur at all positions within the gRNA–TS DNA heteroduplex (**Figure S12B**), in agreement with previous studies (Boyle et al., 2017; Boyle et al., 2021; Doench et al., 2016; Jones et al., 2021; Zhang et al., 2020). This suggests that rA-dA mismatches do not encounter steric barriers within Cas9 that would disfavour their presence, which is consistent with the absence of specific contacts with Cas9 along the length of the major groove of the heteroduplex (**Figure S6**).

To gain further insights into the mechanism of rA-dA mismatch accommodation and extrapolate our structural observations to other heteroduplex positions, we performed molecular dynamics (MD) simulations by introducing single rA-dA mismatches at all 20 positions along the heteroduplex in the context of the catalytically active state of Cas9 (PDB: 6O0Y) and compared the resulting trajectories to the corresponding on-target system. rA-dA mismatch simulations at positions 5, 18, and 19 showed very good agreement with the determined crystal structures (*PTPRC-tgt2 off-target #1*, *FANCF off-target #6*, *AAVS1 off-target #2)*, predicting both dA base extrusion and phosphate backbone distortions (**Figure S12C-D**). In-depth analysis of the rA-dA conformational dynamics was performed by computing the geometrical descriptors defining the base pair complementarity along the heteroduplex (**Figure S12D, Figure S13**) (Lavery et al., 2009). These reveal that at all positions, the rA-dA mismatch is primarily accommodated by base extrusion from the duplex, with positions 3, 5, 15, 16, 18, 19 and 20 undergoing dA base extrusion, and positions 10 and 17 rA base extrusion. Notably, at duplex positions 2, 4, 6, 8, 11 and 12, the broad or bimodal distributions of base pair shear and opening parameters are indicative of considerable conformational dynamics, allowing the rA-dA mismatch to be accommodated by either rA or dA base extrusion (**Figure S13A**).

### PAM-proximal mismatches are accommodated by TS distortion due to seed sequence rigidity

The seed sequence of the guide RNA (nucleotides 11-20) makes extensive interactions with Cas9, both in the absence and presence of bound DNA (Anders et al., 2014; Jiang et al., 2015; Nishimasu et al., 2014; Zhu et al., 2019). Structural pre-ordering of the seed sequence by Cas9 facilitates target DNA binding and contributes to the specificity of on-target DNA recognition (Jiang et al., 2015; O’Geen et al., 2015; Wu et al., 2014). Conversely, binding of off-target DNAs containing PAM-proximal mismatches is inhibited and results in accelerated off-target dissociation (Boyle et al., 2017; Boyle et al., 2021; Ivanov et al., 2020; Jones et al., 2021; Singh et al., 2016; Zhang et al., 2020). Nevertheless, Cas9 does tolerate most base mismatch types within the seed region of the guide RNA, leading to detectable off-target DNA cleavage (Boyle et al., 2021; Doench et al., 2016; Jones et al., 2021; Zhang et al., 2020). In particular, the first two PAM-proximal positions display a markedly higher tolerance for mismatches than the rest of the seed region (Cofsky et al., 2022; Doench et al., 2016; Hsu et al., 2013; Mekler et al., 2017; Zeng et al., 2018); this is supported by our SITE-Seq assay data as the frequency of mismatches at PAM-proximal positions 18-20 is 1.4-2.0x higher than at seed positions 14-17 (**Figure S1D-E**).

Unlike the seed region of the guide RNA, the complementary PAM-proximal TS nucleotides are not directly contacted by Cas9 in the pre-cleavage state and are thus under fewer steric constraints (**Figure S6**), with the exception of duplex position 20 in which the phosphodiester group of the TS nucleotide makes extensive interactions with the phosphate lock loop of Cas9 (Anders et al., 2014) (**Figure S14A**). In agreement with this, four of our off-target complex structures reveal that PAM-proximal base mismatches are accommodated solely by structural distortions of the TS backbone (**Figure 4**), while the conformation of the guide RNA backbone and base stacking within the seed region remain unperturbed (**Figure S14B-C)**. The presence of an rA-dA mismatch in the PAM-proximal position 18 results in the extrusion of the TS nucleobase into the major groove (**Figure 4A**), likely due to steric constraints on duplex width at this position. In contrast, the rA-dA mismatch at duplex position 19 is instead accommodated by a marked distortion in the TS backbone that results in increased duplex width, which preserves base stacking within the duplex in the absence of productive pairing between the adenine bases (**Figure 4B**, **Figure S14B**). Similarly, the rA-dG mismatch at position 19 is accommodated by a ~2 Å displacement of the TS backbone, increasing duplex width. This not only preserves base stacking but also facilitates rA-dG base paring by two hydrogen bonding interactions via their Watson-Crick edges (**Figure 4C**, **Figure S14C**). This off-target complex also contains a rU-dG mismatch at duplex position 20 which does not undergo wobble base pairing as the rU20 nucleotide is extensively contacted by Cas9 (**Figure S6**) and unable to shift towards the major groove and is instead accommodated by a slight shift in the dG nucleotide (**Figure 4C**). Finally, the rU-dT base mismatch at duplex position 20 in the *AAVS1* off-target #4 complex remains unpaired and the dT base lacks ordered electron density (**Figure 4D**). This is likely a result of the dT nucleotide maintaining contact with the phosphate-lock loop of Cas9 (**Figure S14A**), which prevents a reduction in the duplex width and precludes productive base pairing. Overall, these observations indicate that off-target DNAs containing mismatches to the seed sequence of the guide RNA can be accommodated by Cas9 due to limited interactions with the TS DNA in the seed-binding region

**Figure 4.**
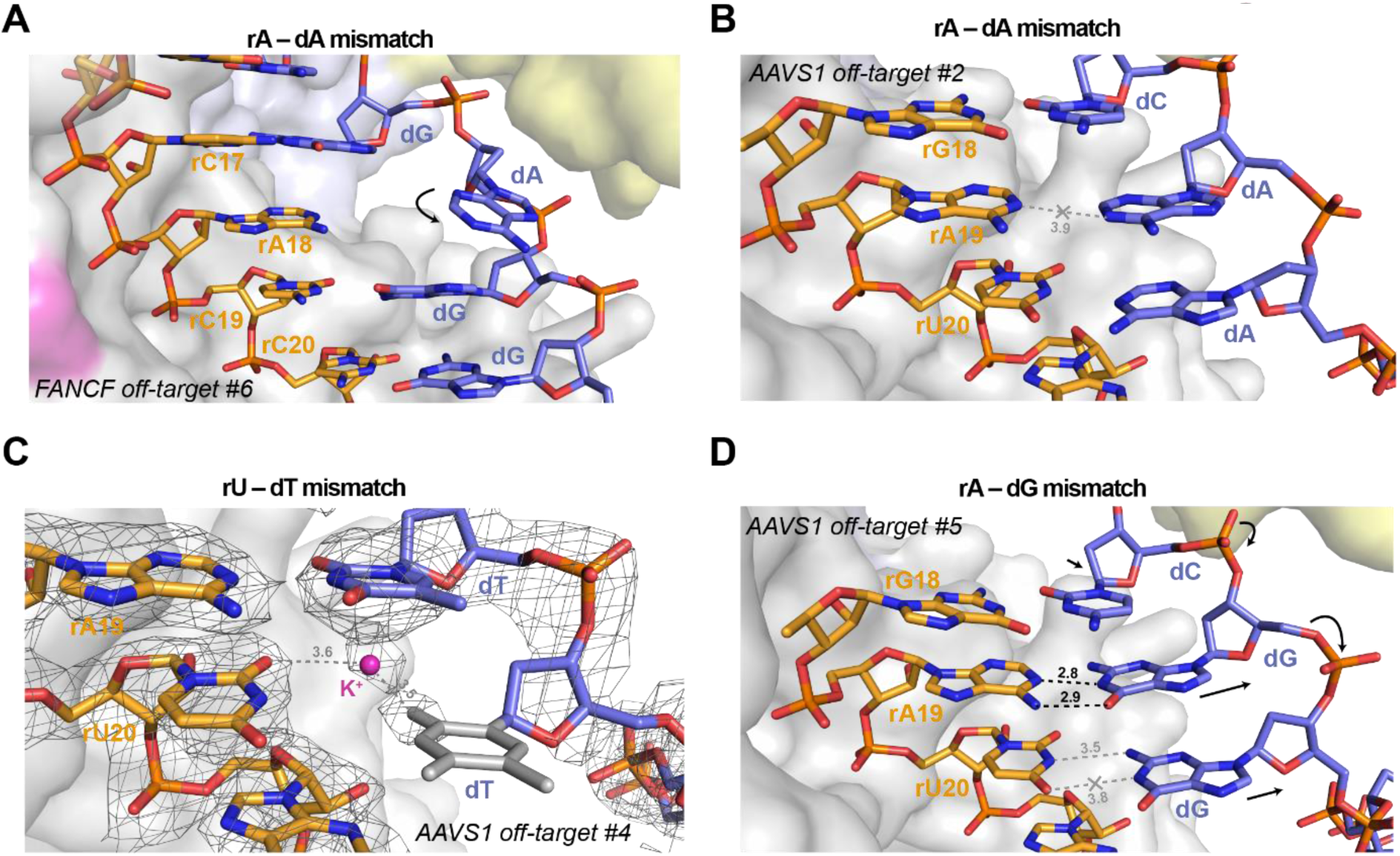
TS distortion facilitates mismatch accommodation in the seed region of the guide–off-target heteroduplex. (**A**) Close-up view of the rA-dA mismatch at position 18 in *FANCF* off-target #6 complex, showing major groove extrusion of the dA base. (**B**) Close-up view of the rA-dA mismatch at position 19 in *AAVS1* off-target #2 complex, showing retention of the dA base in the duplex stack. (**C**) Zoomed-in view of the rA-dG base pair at position 19 and the unpaired rU-dG mismatch at position 20 in the *AAVS1* off-target #5 complex. (**D**) Close-up rU-dT mismatch at the PAM-proximal position 20 in *AAVS1* off-target #4 complex. Residual electron density indicates the presence of an ion or solvent molecule. Refined 2m*F*_o_−D*F*_c_ electron density map of the heteroduplex, contoured at 1.5σ, is rendered as a grey mesh. Structurally disordered thymine nucleobase for which no unambiguous density is present is in grey. Arrows indicate conformational changes in the TS backbone relative to the on-target complex.

### Cas9 recognizes off-targets with single-nucleotide deletions by base skipping or via multiple mismatches

A substantial fraction of off-target sites recovered in our SITE-Seq assay analysis (46.4%, when not considering the possibility of nucleotide insertions or deletions) contained six or more mismatched bases to the guide RNA (**Figure S1C**, **Table S1, Table S2**). Such off-target sequences have previously been proposed to be accommodated by bulging out or skipping of nucleotides (Boyle et al., 2021; Cameron et al., 2017; Doench et al., 2016; Jones et al., 2021; Lin et al., 2014; Tsai et al., 2015), which would result in a shift of the nucleotide register to re-establish correct base pairing downstream of the initially encountered mismatch. The *PTPRC-tgt2* off-target #1 and *FANCF* off-target #3 sites are predicted to contain single nucleotide deletions at duplex positions 15 and 17, respectively (**Figure 1A**). The structures of the corresponding off-target complexes reveal that the single nucleotide deletions in these off-target substrates are not accommodated by bulging out the unpaired guide RNA nucleotide. Instead, the conformations of the guide RNAs remain largely unperturbed and the off-target TS DNAs “skip over” the unpaired gRNA bases to resume productive base-pairing downstream (**Figure 5A-B**). The seed regions of the gRNAs make extensive interactions with the bridge helix and the REC1 domain, whereas the TS DNA phosphate backbones are displaced by almost 3 Å (**Figure S17A-B**). The base pair skips are accommodated by considerable buckling and tilting of the base pairs immediately downstream of the skip site (**Table S6**). An additional consequence of the base pairing register shift is the formation of non-canonical base pairs between the off-target DNA and the extra 5’-terminal guanine nucleotides present in the guide RNA as a consequence of *in vitro* transcription by T7 RNA polymerase (**Figure S17C-D**). This potentially explains the impact of the 5’-guanines on both R-loop stability and *in vitro* cleavage activity (Kulcsar et al., 2020; Mullally et al., 2020; Okafor et al., 2019).

**Figure 5.**
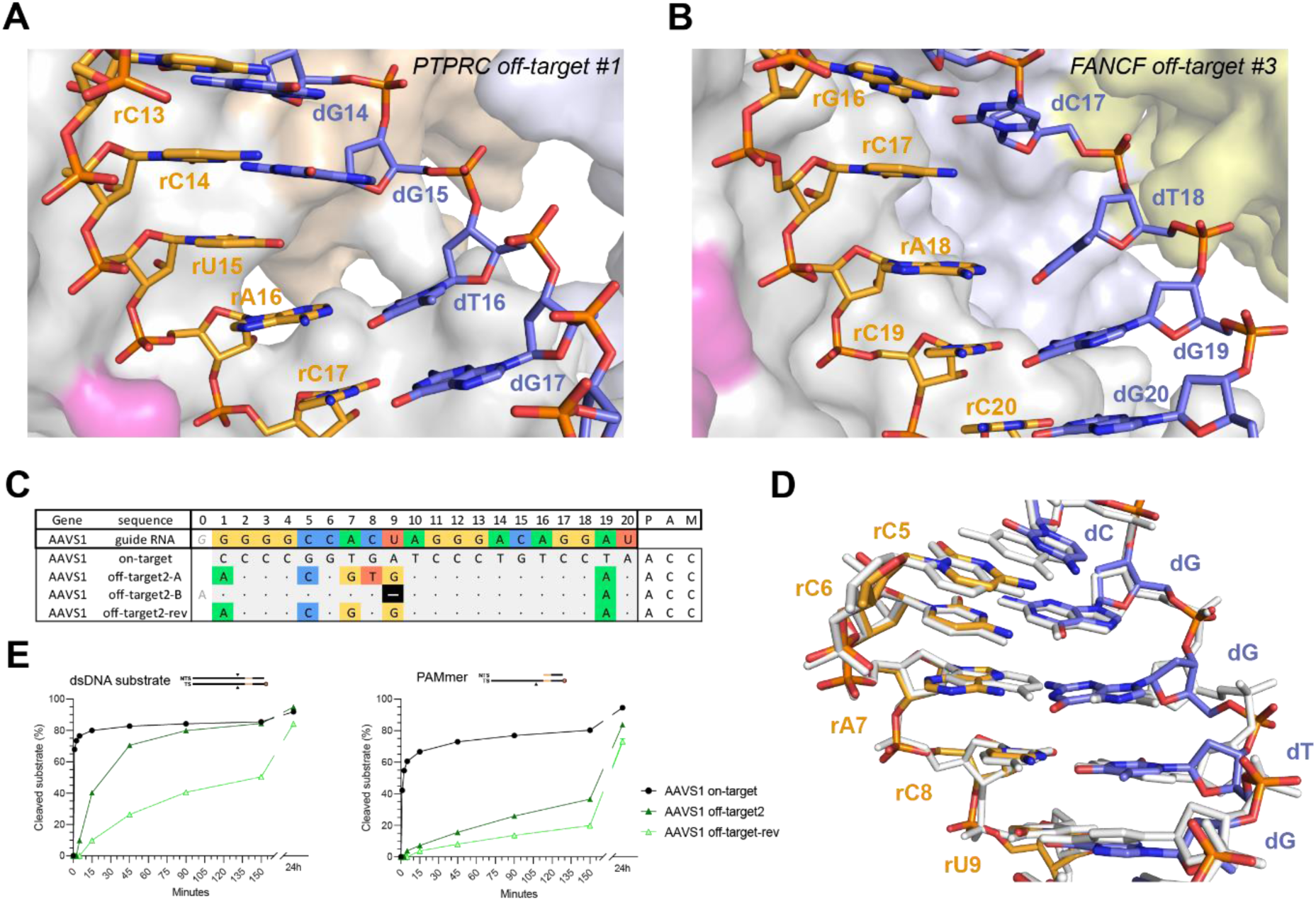
Off-targets with single-nucleotide deletions are accommodated by base skipping or multiple consecutive mismatches. (**A**) Zoomed-in view of the base skip at duplex position 15 in the *PTPRC-tgt2* off-target #1 complex. (**B**) Zoomed-in view of the base skip at duplex position 17 in the *FANCF* off-target #3 complex. (**C**) Schematic depiction of alternative base pairing interactions in the *AAVS1* off-target #2 complex. *AAVS1* off-target #2-rev substrate was designed based on the *AAVS1* off-target #2, with the reversal of a single mismatch in the consecutive region back to the corresponding canonical base pair. (**D**) Structural overlay of the *AAVS1* off-target #2 (coloured) and *AAVS1* on-target (white) heteroduplexes. (**E**) Cleavage DNA kinetics of AAVS1 on-target, off-target #2 and off-target #2-rev substrates.

Originally, our SITE-Seq assay analysis classified the *AAVS1* off-target #2 as a single-nucleotide deletion at duplex position 9 (**Figure 5C**). Unexpectedly, the structure of the *AAVS1* off-target #2 complex instead reveals that the off-target substrate is bound in the unshifted register, resulting in the formation of five base mismatches in the PAM-distal half of the guide RNA–TS duplex (**Figure 5D**), including a partially paired rC-dC mismatch at position 5, an rA-dG Hoogsteen pair at position 7, a partially paired rC-dT mismatch at position 8, and a tautomeric rU-dG pair at position 9. The backbone conformations of the gRNA and the off-target TS exhibit minimal distortions and are nearly identical with the corresponding on-target heteroduplex (**Figure 5D**). This implies that some mismatch combinations might synergistically result in guide RNA and TS backbone geometries that mimic the on-target conformation and result in off-target tolerance. To test this hypothesis, we reverted the rC-dT mismatch at position 8 to the on-target rC-dG pair, thereby reducing the total amount of off-target mismatches from 6 to 5 (**Figure 5C**). The resulting off-target substrate (*AAVS1* off-target #2-rev) exhibited substantially reduced cleavage rates in both dsDNA and PAMmer formats, as well as a significantly increased dissociation rate (**Figure 5E, Table S8**). We replicated these results for two other *AAVS1* off-target substrates with different mismatch patterns that were initially predicted to contain PAM-distal deletions (**Figure S18, Table S8**). Together, these results suggest that some *bona fide* off-target substrates containing multiple consecutive mismatches, the reversal of one mismatch may affect the structural integrity of the guide RNA– TS DNA heteroduplex and interfere with DNA binding and/or conformational activation of Cas9, despite a reduction in the total number of mismatches.

To further investigate the accommodation of PAM-distal deletions in Cas9 off-target sites, we selected an additional efficiently cleaved off-target substrate (*CD34* off-target #9, **Figure S19A,B**), containing a dT substitution at position 17 (resulting in a rU-dT mismatch with the *CD34* sgRNA) and a predicted single-nucleotide deletion at position 6. Crystal structure of the resulting complex revealed that the rU-dT mismatch involves the *syn* conformer of the rU17 nucleotide (**Figure S19C**). The deletion at position 6 is accommodated by base skipping, leaving the base of rA6 unpaired within the guide-TS heteroduplex stack (**Figure S19E**) and the register shift results enables the 5’-terminal guanine nucleotide of the gRNA (introduced during *in vitro* transcription) to base pair with 3’-terminal TS nucleotide (**Figure S19F**). The backbone distortions in the TS are accompanied by compensatory rearrangements of the REC2 and REC3 domains of Cas9 (**Figure S19G**), which preserve all contacts with the heteroduplex (**Figure S6**).

Collectively, these results indicate that deletion-containing off-target complexes are accommodated either by RNA base skipping, as opposed to RNA nucleotide bulging, or by the formation of multiple base mismatches, with the precise mechanism dependent in part on the position of the deletion.

### PAM-distal mismatches perturb the Cas9 conformational checkpoint

*FANCF* off-target #4, which contains three PAM-distal mismatches at positions 1-3 and a G-U mismatch in position 10 (**Figure 1A**), is reproducibly the top ranking off-target site for the *FANCF* guide RNA, as detected by SITE-Seq assay analysis at the lowest Cas9 RNP concentrations (**Table S1, Table S2**). The off-target substrate exhibits slow cleavage kinetics *in vitro* with both dsDNA and PAMmer substrates (**Figure 2B, Figure S2A-B**), indicating a perturbation of the conformational activation checkpoint of Cas9. The structure of the *FANCF* off-target #4 complex reveals that the RNA–DNA heteroduplex is unpaired at positions 1-3 as a result of the PAM-distal mismatches, with nucleotides 1-2 of the guide RNA and 19-20 of the TS disordered (**Figure 6A**). Furthermore, Cas9 undergoes structural rearrangements of its REC lobe and the HNH domain (**Figure 6B**), resulting in a root mean square displacement of the REC2 and REC3 domains of 3.7 Å (1,315 Cα atoms) relative to the *FANCF* on-target complex structure. The REC3 domain undergoes a 19-degree rotation (**Figure 6B**), facilitated by extending the helix comprising residues 703-712 through restructuring of loop residues 713-716 (**Figure S20A**), to accommodate the altered guide RNA conformation. The REC2 domain rotates 32° away from the REC3 domain (**Figure 6B**). This is accompanied by restructuring of the hinge loop residues 174-180 and disordering of loops 258-264, 284-285, and 307-309. Concomitantly, the HNH domain rotates 11° away from the heteroduplex, as compared to the *FANCF* on-target structure, to accommodate distortion of the TS DNA (**Figure 6B**).

**Figure 6.**
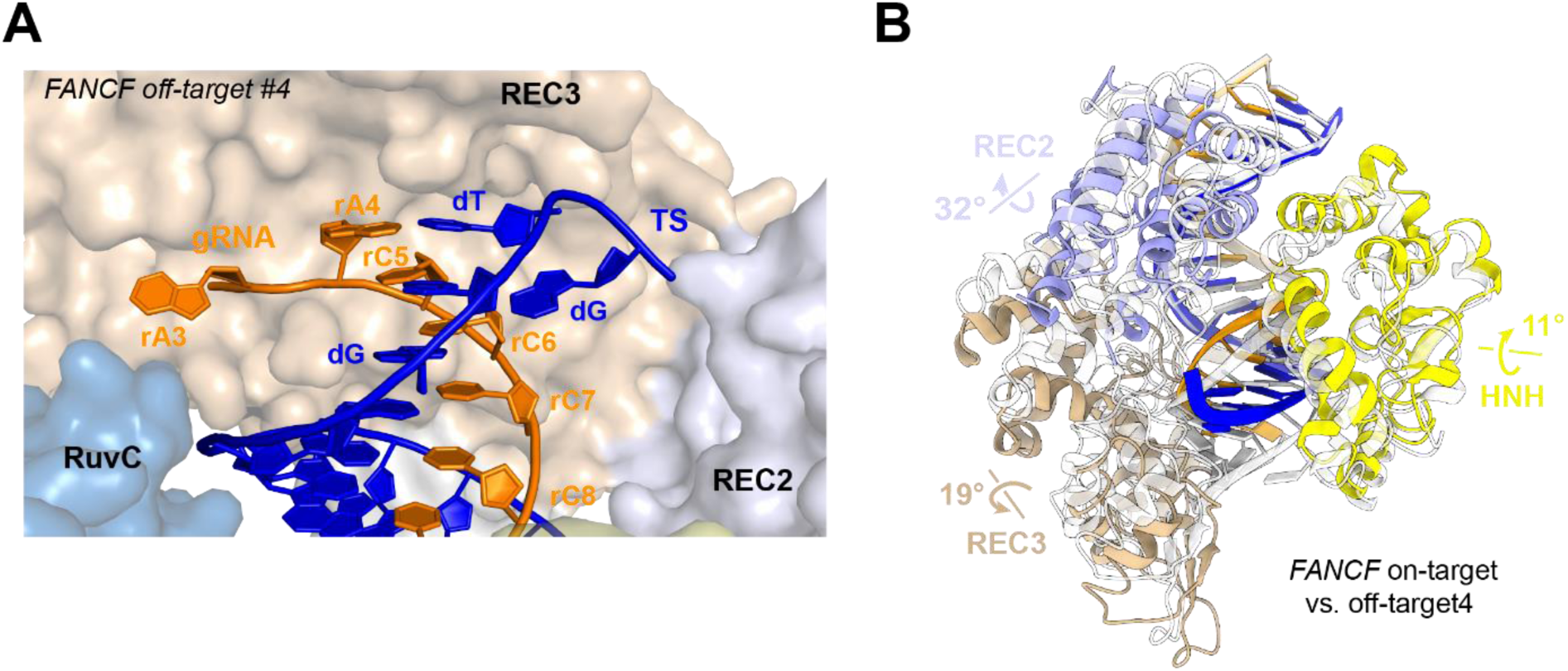
*FANCF* off-target #4 exhibits conformational changes in the REC2/3 and HNH domains due to PAM-distal duplex unpairing. (**A**) Close-up view of the unpairing of mismatched bases at the PAM-distal end of the *FANCF* off-target #4 heteroduplex. The last two nucleotides on each strand could not be modelled due to structural disorder. (**B**) Overlay of the *FANCF* off-target #4 and *FANCF* on-target complex structures. The *FANCF* off-target #4 complex is coloured according to the domain legend in Figure 1A, *FANCF* on-target complex is shown in white. The REC1, RuvC, and PAM-interaction domains have been omitted for clarity, as no structural differences are observed.

The unpaired 5’ end of the sgRNA is located at the interface between the REC3 and the RuvC domain and maintains interactions with heteroduplex-sensing residues Lys510, Tyr515, and Arg661 of the REC3 domain (**Figure S20B**). In contrast to the corresponding on-target complex structure, the unpaired 3’ end of the off-target TS breaks away from the REC3 lobe and instead points towards the REC2 domain, forming unique interactions with Arg895, Asn899, Arg905, Arg919 and His930 in the HNH domain (**Figure S20C**). These interactions (**Figure S20D**) could be responsible for the observed repositioning of the REC lobe and HNH domain, and they may impede the formation of a cleavage-competent complex.

The conformation of the *FANCF* off-target #4 complex is distinct from the conformations observed in cryo-EM reconstructions of the pre- and post-cleavage states of the Cas9 complex (Zhu et al., 2019) (**Figure S21A-B**). Instead, the off-target complex structure most closely resembles that of a high-fidelity variant xCas9 3.7 containing amino acid substitutions that disrupt interactions with the TS DNA (Guo et al., 2019). Although the xCas9 3.7 complex adopts a slightly different REC lobe conformation (**Figure S21C**), the PAM-distal duplex also undergoes unpairing at positions 1–3 and displays a comparable degree of structural disorder (**Figure S21D**). These structural observations thus suggest that the presence of multiple mismatches in the PAM-distal region of a guide RNA–off-target DNA duplex leads to conformational perturbations in the DNA-bound complex that resemble the structural consequences of specificity-enhancing mutations in high-fidelity Cas9 variants.

## DISCUSSION

The off-target activity of Cas9 has been extensively documented in prior genome editing, biochemical and biophysical studies (Boyle et al., 2017; Boyle et al., 2021; Doench et al., 2016; Jones et al., 2021; Lazzarotto et al., 2020; Zhang et al., 2020). Although numerous methods have been devised for computational prediction of genomic off-target sites and their experimental validation, these have reported highly variable mismatch tolerance profiles depending on the screening method and the target sequence. Thus, a comprehensive understanding of this phenomenon is still lacking, particularly as to whether off-target tolerance has an underlying structural basis. In this study, we used the SITE-Seq assay to examine the off-target landscape of 12 well-studied guide RNAs, observing a broad variation of cleavage activities associated with individual off-target substrates. To shed light on the molecular mechanisms underpinning off-target activity, we determined atomic structures of a representative set of 16 off-target complexes, thus providing fundamental insights into the structural aspects of off-target recognition.

### Role of non-canonical base pairing in off-target recognition

The principal, and largely unexpected, finding of our structural analysis is that the majority of nucleotide mismatches in *bona fide* off-target substrates are accommodated by non-canonical base pairing interactions. These range from simple rG-dT/rU-dG wobble or rA-dG/rG-dA/rG-dG Hoogsteen base pairing interactions, to pyrimidine-pyrimidine pairs that rely on (deoxy)ribose-phosphate backbone distortions that reduce duplex width. With the notable exception of rA-dA mismatches, which are accommodated by extrahelical base extrusion, the structural rearrangements associated with base mismatch accommodation largely preserve base stacking, which is the primary determinant of nucleic acid duplex stability (Yakovchuk et al., 2006). For some off-target sequences, our structures are suggestive of base protonation or tautomerization facilitating hydrogen bonding interactions in otherwise non-permissive mismatches, such as rA-dC. These rare base pair forms have been previously observed in both RNA and DNA duplexes and are thought to be important contributors to DNA replication and translation errors (Kimsey et al., 2015; Kimsey et al., 2018). Future studies employing complementary structural methods, such as nuclear magnetic resonance, will help confirm the occurrence of non-canonical base states in off-target complexes.

The mismatch tolerance of Cas9 can be explained primarily by two factors. Firstly, Cas9 does not directly contact the major- or minor-groove edges of the guide RNA–TS DNA heteroduplex base pairs at any of the duplex positions and thus lacks a steric mechanism to enforce Watson-Crick base pairing (**Figure S6**). This is further underscored by Cas9’s tolerance of base modifications in target DNA, including cytosine 5-hydroxymethylation and, at least at some duplex positions, glucosyl-5-hydroxymethylation (Vlot et al., 2018). In this respect, Cas9 differs from other molecular systems, notably the ribosome and replicative DNA polymerases, which enhance the specificity of base-pairing by direct readout of base-pair shape and steric rejection of mismatches (Kunkel and Bebenek, 2000; Rodnina and Wintermeyer, 2001; Timsit, 1999).

Secondly, Cas9 is a multidomain protein that displays considerable conformational plasticity and is therefore able to accommodate local distortions in the guide–TS duplex geometry by compensatory rearrangements of the REC2, REC3 and HNH domains (Chen et al., 2017; Donohoue et al., 2021). Indeed, in most off-target structures reported in this study, almost all atomic contacts between Cas9 and the guide–TS heteroduplex are preserved (**Figure S6**). Thus, Cas9 only detects guide-target mismatches by indirect readout of the guide RNA– TS DNA heteroduplex width, except at the PAM-distal end of the heteroduplex where base mismatches result in duplex unpairing, as discussed below. Our observations are consistent with recent molecular dynamics simulation studies showing that internally positioned mismatches within the guide RNA–TS DNA heteroduplex are readily incorporated within the heteroduplex and have only minor effects on Cas9 interactions (Mitchell et al., 2020). The lack of a steric base-pair enforcement mechanism and the resulting off-target promiscuity likely reflects the biological function of Cas9 in CRISPR immunity, where mismatch tolerance enables targeting of closely related viruses and hindering immune evasion by mutations or covalent base modifications (Deveau et al., 2008; Semenova et al., 2011; van Houte et al., 2016; Yaung et al., 2014). On the other hand, such conformational plasticity also enables the incorporation of various chemical modifications in the guide RNA that are compatible with the A-form geometry of the heteroduplex (Cromwell et al., 2018; Donohoue et al., 2021; Hendel et al., 2015; O’Reilly et al., 2019; Yin et al., 2018; Yin et al., 2017). These could potentially disfavour the formation of certain non-canonical base pairs or reduce the backbone flexibility of the guide RNA, thereby enhancing off-target discrimination. However, the effect of specific guide RNA modifications on particular types of mismatches is yet to be closely examined mechanistically (Donohoue et al., 2021).

### Structural rigidity of the guide RNA seed region and implications for off-target recognition

The seed sequence of the Cas9 guide RNA (nucleotides 11-20) is the primary determinant of target DNA binding, a consequence of its structural pre-ordering in an A-like conformation by extensive interactions with Cas9 (Anders et al., 2014; Jiang et al., 2015; Nishimasu et al., 2014; Zhu et al., 2019). Our data indicate that structural rigidity of the guide RNA seed sequence also affects off-target recognition, as base mismatches in the seed region of the heteroduplex can only be accommodated by conformational distortions of TS DNA, which is subject to only a few steric constraints, notably at position 20 due to interactions with the phosphate lock loop (Anders et al., 2014). This increases the energetic penalty of base mispairing in the seed region of the heteroduplex, and thus contributes to mismatch sensitivity of Cas9 within the seed region. Although structural distortions of TS DNA facilitate binding of off-target substrates containing seed mismatches, they may nevertheless inhibit off-target cleavage by steric hindrance of the HNH domain, thereby further contributing to off-target discrimination. The contrasting structural plasticities of the guide RNA and TS DNA strands are manifested in the differential activities of Cas9 against off-targets containing rU-dG and rG-dT mismatches within the seed region (Boyle et al., 2021; Doench et al., 2016; Hsu et al., 2013; Jones et al., 2021; Zhang et al., 2020). Whereas rG-dT mismatches can be readily accommodated by wobble base pairing, seed sequence rigidity is expected to hinder rU-dG wobble base pairing. Combined with a lower energetic penalty associated with rG-dT mismatch binding (binding an off-target with an rG-dT mismatch requires unpairing a dT-dA base pair in the off-target DNA, while rU-dG off-target recognition requires dC-dG unpairing), these effects thus help Cas9 discriminate against rU-dG mismatches in the seed region.

### Recognition of off-targets containing insertions and deletions

*Bona fide* off-target sites containing insertions or deletions have been detected in a number of studies (Boyle et al., 2021; Cameron et al., 2017; Doench et al., 2016; Jones et al., 2021; Tsai et al., 2015). Nucleotide “bulging” has been proposed as a mechanism to recognize such off-targets, which would otherwise result in large numbers of consecutive base mismatches. However, as Cas9 encloses the guide RNA–TS DNA heteroduplex in a central channel and makes extensive interactions along the entire length of the guide RNA strand, the formation of RNA bulges is precluded by steric clashes, pointing to a different mechanism.

Indeed, our structural analysis provides no evidence for RNA bulging. Instead, the structures of *PTPRC-tgt2* off-target #1 and *FANCF* off-target #3 complexes reveal that off-target sequences predicted to contain single-nucleotide deletions in the seed region of the heteroduplex are recognized by base skipping, resulting in an unpaired guide RNA base within the duplex stack. This is enabled by the lack of extensive contacts of Cas9 with the TS, while rigid coordination of the guide RNA in the seed region disfavours extrahelical guide RNA bulging. However, base skipping in the seed region of the heteroduplex likely incurs a large energetic penalty. As the seed region of the TS DNA is devoid of Cas9 contacts in the pre-cleavage state (Zhu et al., 2019), off-targets containing single-nucleotide insertions in the seed region of the heteroduplex are likely to be recognized by DNA nucleotide bulging, likewise incurring a large energetic penalty as unwinding an off-target DNA sequence containing an insertion requires breaking an extra base pair. Additionally, TS DNA distortion might inhibit cleavage by steric hindrance of the HNH domain. These observations thus explain why Cas9 appears to tolerate mismatches better than insertions or deletions and why deletions and insertions within the seed region are particularly deleterious (Boyle et al., 2021; Cameron et al., 2017; Doench et al., 2016; Jones et al., 2021).

In contrast, off-target sequences containing insertions or deletions in the PAM-distal region of the heteroduplex (positions 1-10) can be recognized either by base skipping, as in the case of *CD34* off-target #9, or bound in the unshifted register, with multiple base mismatches accommodated by non-canonical base pairing interactions, as seen for *AAVS1* off-target #2, which was previously predicted to contain an RNA bulge or skip (Cameron et al., 2017; Lazzarotto et al., 2020). Which of the two mechanisms is used likely depends on the off-target sequence and the position of the deletion, which in turn dictate the number of mismatches between the guide and the off-target sequence in the unshifted register. Off-target accommodation by multiple non-canonical base pairs likely relies on their synergistic effect to mimic the on-target heteroduplex geometry, which enables unperturbed binding, as supported by our observations that mismatch reversal in several off-target substrates reduces their rates of cleavage. Although our structural analysis does not examine off-target substrates containing insertions, we posit that PAM-distal insertions are recognized as multiple mismatches due to steric hindrance of extrahelical bulging. Thus, our structural findings suggest that a substantial fraction of off-target sites predicted to contain insertions or deletions may be bound via multiple mismatches instead (Boyle et al., 2021; Doench et al., 2016; Jones et al., 2021).

### PAM-distal base pairing and the conformational checkpoint of Cas9

Upon substrate DNA hybridization and R-loop formation, Cas9 undergoes conformational activation of its nuclease domains (Bravo et al., 2022; Pacesa et al., 2022; Zhu et al., 2019). The Cas9 REC3 domain plays a key role in the process, as it senses the integrity of the PAM-distal region of the guide RNA–TS DNA heteroduplex and allosterically regulates the REC2 and HNH domains, providing a conformational checkpoint that traps Cas9 in a conformationally inactive state in the absence of PAM-distal hybridization (Chen et al., 2017; Dagdas et al., 2017; Palermo et al., 2018; Zhu et al., 2019). Our structural data confirm that mismatches at the PAM-distal end of the heteroduplex (positions 1-3) result in heteroduplex unpairing, incomplete R-loop formation and structural repositioning of the REC3 domain (**Figure 6**), indicating a perturbation of the Cas9 conformational checkpoint. We envision that the observed conformational state mimics the structural effect of 5’-truncated guide RNAs, which have been shown to improve targeting specificity (Fu et al., 2014). Furthermore, similarities with the structure of a high-fidelity Cas9 variant (Guo et al., 2019) suggest a shared underlying mechanism for increased specificity.

### Implications for off-target prediction

Our structural data reveal that Cas9 plays a limited steric role in off-target discrimination insofar as only sensing the integrity and general shape of the guide–target heteroduplex. Off-target activity is thus largely determined by the kinetics and energetics of R-loop formation, that is off-target DNA strand separation and concomitant guide RNA–TS DNA hybridization, and subsequent Cas9 conformational activation. We observe on multiple occasions that a given mismatch adopts different conformational arrangements depending on its position along the guide RNA–TS DNA heteroduplex, as further supported by MD simulations of rA-dA mismatches. This poses a challenge for *ab initio* modelling of off-target activity, as biophysical models of off-target binding and cleavage are bound to be of limited accuracy unless they incorporate position-dependent energetic penalties for each base mismatch type and for deletions, as well as position- and base-specific penalties for insertions (Boyle et al., 2021; Jones et al., 2021; Zhang et al., 2020). As illustrated by our study, MD simulations can complement experimental data to provide structural information on specific mismatches in remaining positions within the heteroduplex. Thus, ongoing structural and computational studies, combined with machine learning approaches, will assist in generating complete models for off-target prediction.

Furthermore, as certain off-target sequences that are incompatible with dsDNA cleavage can undergo NTS nicking (Fu et al., 2019; Jones et al., 2021; Murugan et al., 2020; Zeng et al., 2018), future bioinformatic models need to be able to predict off-target nicking activity as well. Furthermore, accurate modelling of off-target interactions remains difficult due to context-dependent effects, as documented in previous studies showing that the binding and cleavage defects of consecutive mismatches deviate from additivity (Boyle et al., 2021; Cameron et al., 2017; Lazzarotto et al., 2020; Zhang et al., 2020). Indeed, our structural data rationalize this by showing that the conformation of a given base mismatch is highly sensitive to the presence of neighbouring mismatches. As seen in the case of *AAVS1* off-target #2 complex, multiple mismatched bases can synergistically combine to preserve an on-target-like heteroduplex conformation that passes the REC3 conformational checkpoint, supporting nearly on-target efficiencies of cleavage (Zhang et al., 2020). This is in line with recent cryo-EM structural studies suggesting that indirect readout of heteroduplex conformation is coupled to nuclease activation, while mismatches disrupt this coupling (Bravo et al., 2022; Pacesa et al., 2022). Critically, reversion of one of the mismatches in this off-target substrate impairs cleavage activity. Similar effects have been described for other DNA binding proteins such as transcription factors, where mismatches modulate transcription factor binding activity by affecting the conformation of the DNA duplex (Afek et al., 2020). In an analogy with Cas9, these proteins check for correct binding sites through indirect sequence readout by sampling for the correct duplex shape rather than base sequence (Abe et al., 2015; Kitayner et al., 2010; Rohs et al., 2009a; Rohs et al., 2009b).

In conclusion, structural insights presented in this study establish an initial framework for understanding the molecular basis for the off-target activity of Cas9. In conjunction with ongoing computational studies, these findings will help achieve improved energetic parametrization of off-target mismatches and deletions/insertions, thus contributing to the development of more accurate off-target prediction algorithms and more specific guide RNA designs. In doing so, these studies will contribute towards increasing the precision of CRISPR-Cas9 genome editing and the safety of its therapeutic applications.

### Limitations of the study

The structural dataset presented in this study is necessarily restricted in scope. Although all mismatch types are covered, a much larger collection of off-target complex structures would be required to cover all mismatches at all positions to achieve a complete structural overview of the mismatch tolerance underpinning intrinsic off-target activity of Cas9. Although MD simulations can help fill in the gaps in the experimental data, our findings suggest that mismatch accommodation appears to be sequence context-dependent, thus limiting their predictive power. Although we provide insights into off-target substrates containing single-nucleotide deletions, it is presently difficult to predict which mechanism occurs at a given heteroduplex position. Moreover, we currently lack structural data for off-target substrates containing insertions, which will be addressed in future studies. Finally, not all structurally characterized off-targets (~50%) are efficiently cleaved in cells, despite detectable cleavage *in vitro*, implying that other factors affect Cas9 off-targeting in the context of eukaryotic genomes. Thus, further work combining structural and computational approaches will be needed to accurately predict the off-target activity of Cas9.

### Author contributions

M.P., P.C., P.D.D., and M.J. conceived the study. M.P. purified wild-type Cas9, performed *in vitro* cleavage assays, crystallized ternary Cas9 complexes, solved the structures, and performed structural analysis along with M.J.; A.C. performed switchSENSE binding measurements, under the supervision of F.H.T.A; A.S. and P.R.A. performed molecular dynamics simulations, under the supervision of G.P.; M.J.I. performed the SITE-Seq assay; C-H.L. wrote the computational off-target classification model and P.D.D. and P.C. analysed the output; K.B. purified dCas9, transcribed sgRNAs, and prepared DNA substrates for *in vitro* cleavage assays; M.P., P.C., P.D.D., and M.J. wrote the manuscript.

## Acknowledgements

We thank members of the Jinek laboratory and Caribou Biosciences, Inc. for discussion and critical reading of the manuscript. We thank Christelle Chanez and Michael Schmitz for their assistance with preparing reagents. We thank Kenny B. Jungfer for assistance with crystallographic data collection. We thank John Hawkins and Ilya Finkelstein for providing access to the NucleaSEQ prediction pipeline. We thank Vincent Olieric, Meitian Wang, and Takashi Tomizaki (Swiss Light Source, Paul Scherrer Institute) for assistance with crystallographic data collection and Nena Matscheko from Dynamic Biosensors for support with switchSENSE experiments. We thank the NCCR RNA and Disease for providing infrastructural support. This work was supported by the Swiss National Science Foundation Grant 31003A_182567 (to M.J.), and the National Institutes of Health (Grant No. R01GM141329) and National Science Foundation (Grant No. CHE-1905374) (to G.P.). M.J. is an International Research Scholar of the Howard Hughes Medical Institute and Vallee Scholar of the Bert L & N Kuggie Vallee Foundation.

## SUPPLEMENTAL FIGURES AND TABLES

**Figure S1.**
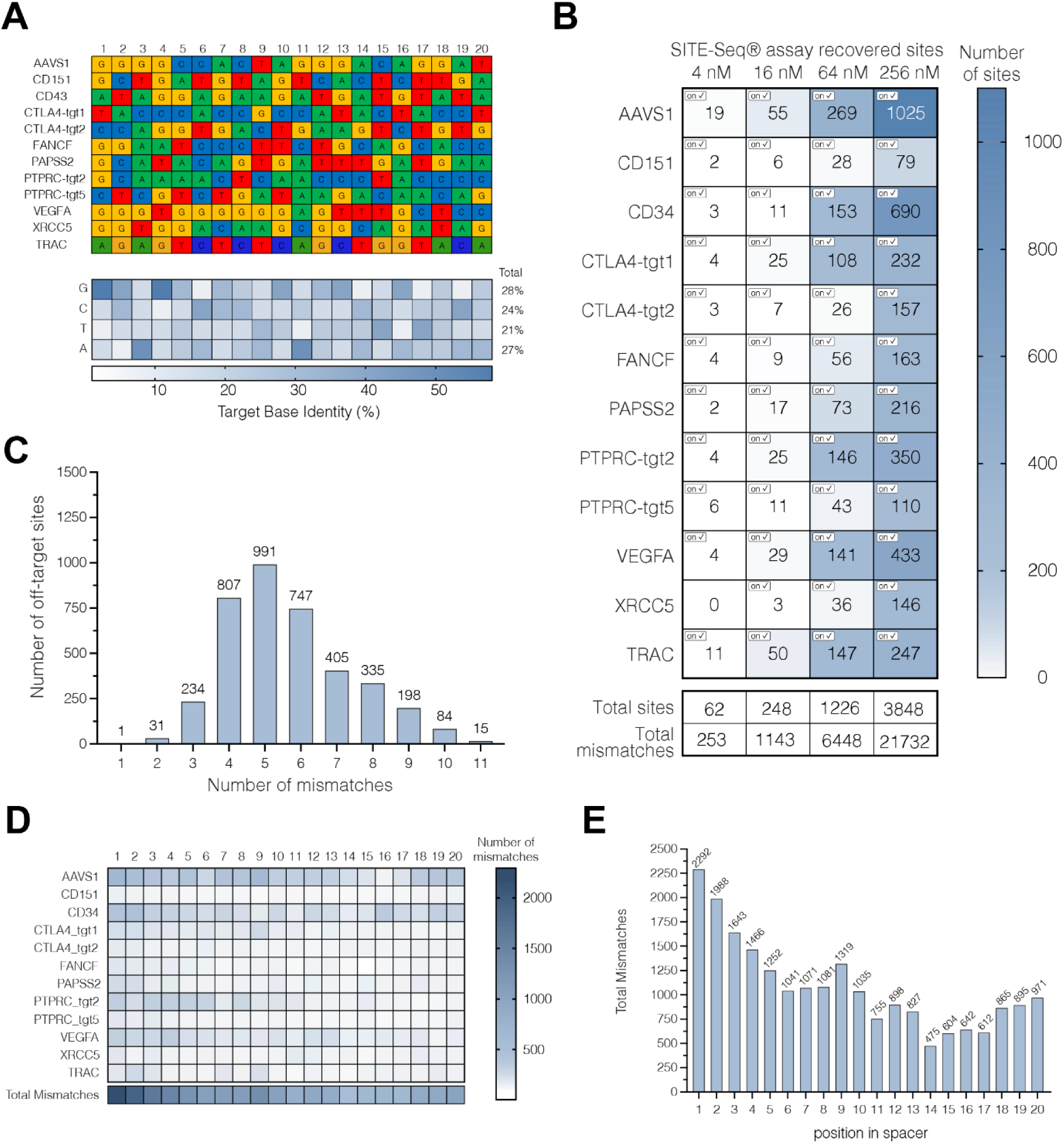
Off-target profiling of selected genomic sites using SITE-Seq. (**A**) Selected genomic targets and the corresponding guide RNA sequences selected for the SITE-Seq assay off-target profiling. Heatmap indicates frequency of nucleotide identity across each position for the selected targets. (**B**) SITE-seq assay analysis for RNPs assembled with indicated crRNAs. The numbers of detected off-target sites are shown as a function of RNP concentration. Checked boxes indicate recovery of the on-target site. n=3 replicates per sample. (**C**) Number of off-target sites recovered by the SITE-Seq assay are shown as a function of the number of mismatches between the guide RNA and the off-target sequence. (**D**) Frequency of nucleotide mismatches at each guide RNA–off-target DNA heteroduplex position for all off-target sites identified in (B). (**E**) Number of total identified mismatches per heteroduplex position.

**Figure S2.**
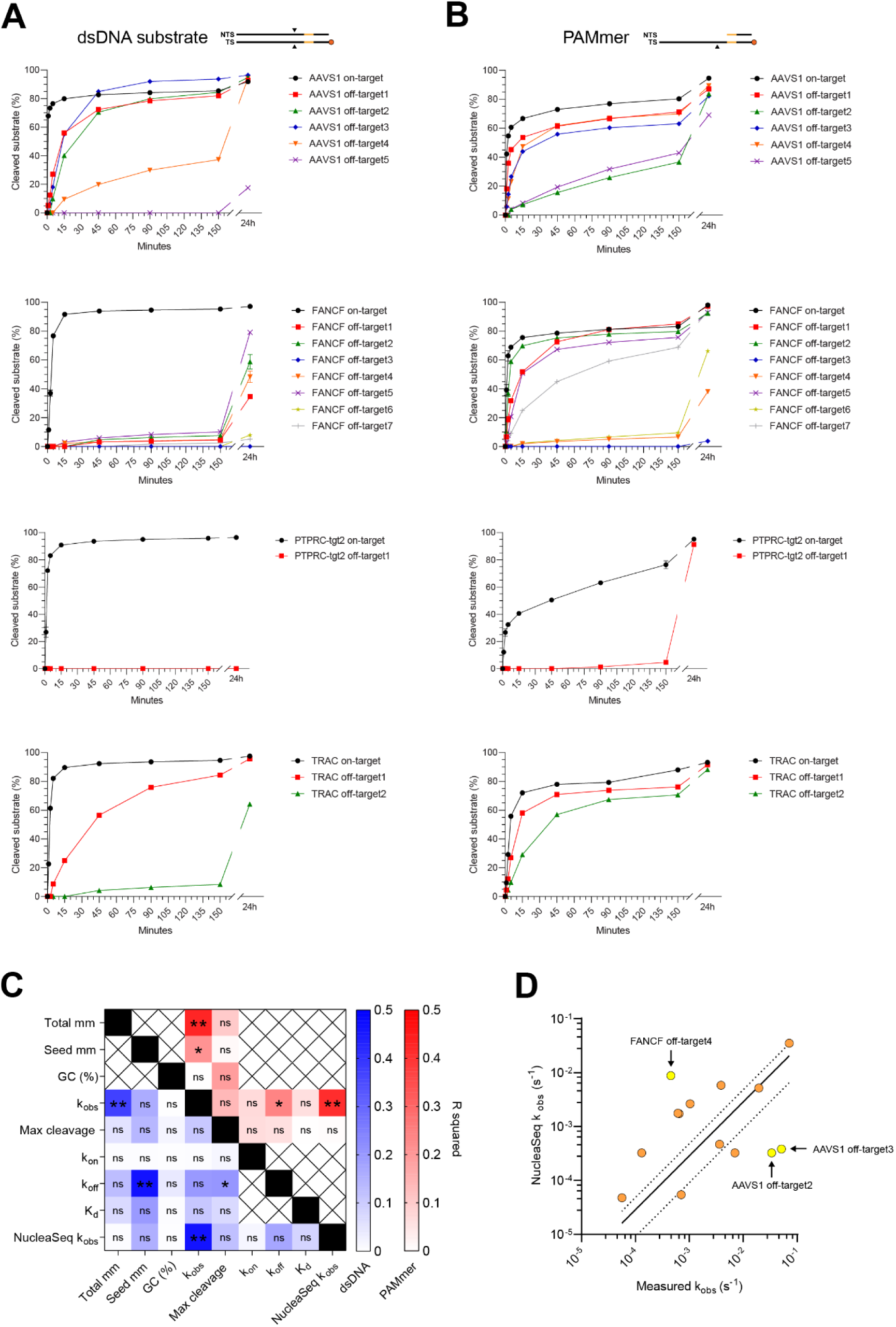
*in vitro* cleavage of a subset of Cas9 off-target substrates. (**A**) *In vitro* clevage kinetics of fully double stranded on- and off-target synthetic DNA substrates labelled on the target-strand for each guide RNA used in the study. Black triangles in the substrate schematic (top) indicate position of cleavage sites. Each data point represents a mean of four independent replicates. Error bars represent standard deviation for each time point. (**B**) *In vitro* clevage kinetics of partially single stranded (PAMmer) on- and off-target substrates. (**C**) Heatmap representation of mutual correlations between measured kinetic and thermodymamic parameters including cleavage (k_obs_), substrate DNA binding (k_on_), substrate dissociation (k_off_) rate constants, equilibrium dissociation constant (K_d_) with numbers of nucleotide mismatches in the off-target sites (total and within seed), the GC content of the spacer (%GC) and cleavage rate predicted using the NucleaSeq algorithm (NucleaSeq k_obs_). The values were calculated across all off-targets for both dsDNA (lower left half, in blue), and partially single stranded (PAMmer) substrates (upper right half, in red). ns, no significant correlation. (**D**) Correlation between measured and NucleaSeq-predicted k_obs_ rate constants. Off-target sites with significant deviations are highlighted in yellow.

**Figure S3.**
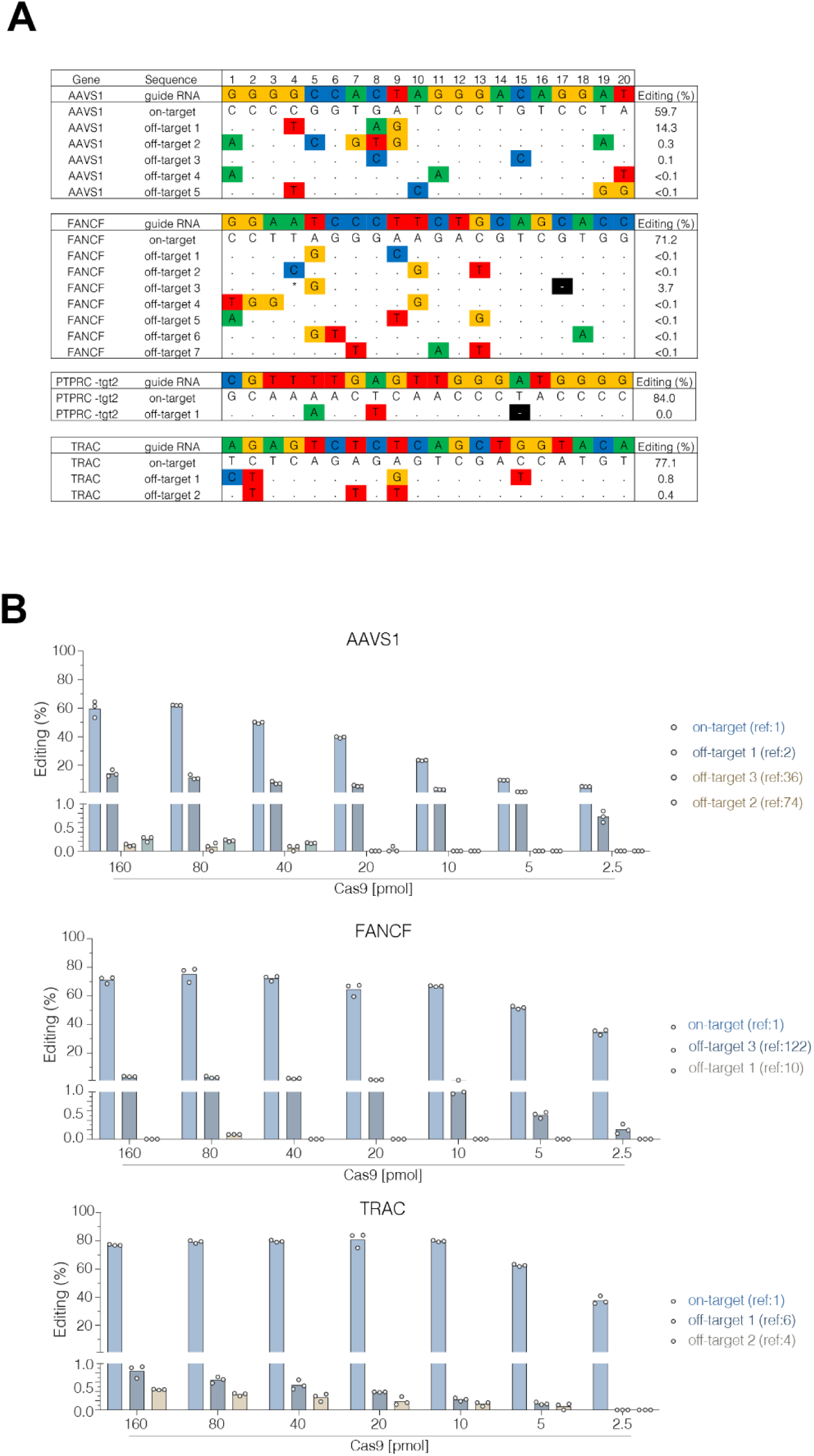
In vivo editing at on- and off-target sites. **(A)** Editing activity of crystalized on- and off-targets in primary human T cells. Matching bases in off-targets are denoted by a dot, whereas base pair mismatches and deletions (–) are highlighted. Asterisk indicated single-nucleotide polymorphism in PBMC donor sequence different from hg38 genomic reference. Editing rates are from highest Cas9 concentration shown in (B), n=3 replicates per sample, limit of detection = 0.1%. **(B)** Dose titration of Cas9 in human T cells for selected on and off-target sites. Off-target numbering correspond to sample shown in **(A)**, and/or reference number (“ref”) as indicated in Table S1. Cas9 concentration shown below x-axis (1:3 molar ratio of Cas9 to guides), N=3 replicates per sample.

**Figure S4.**
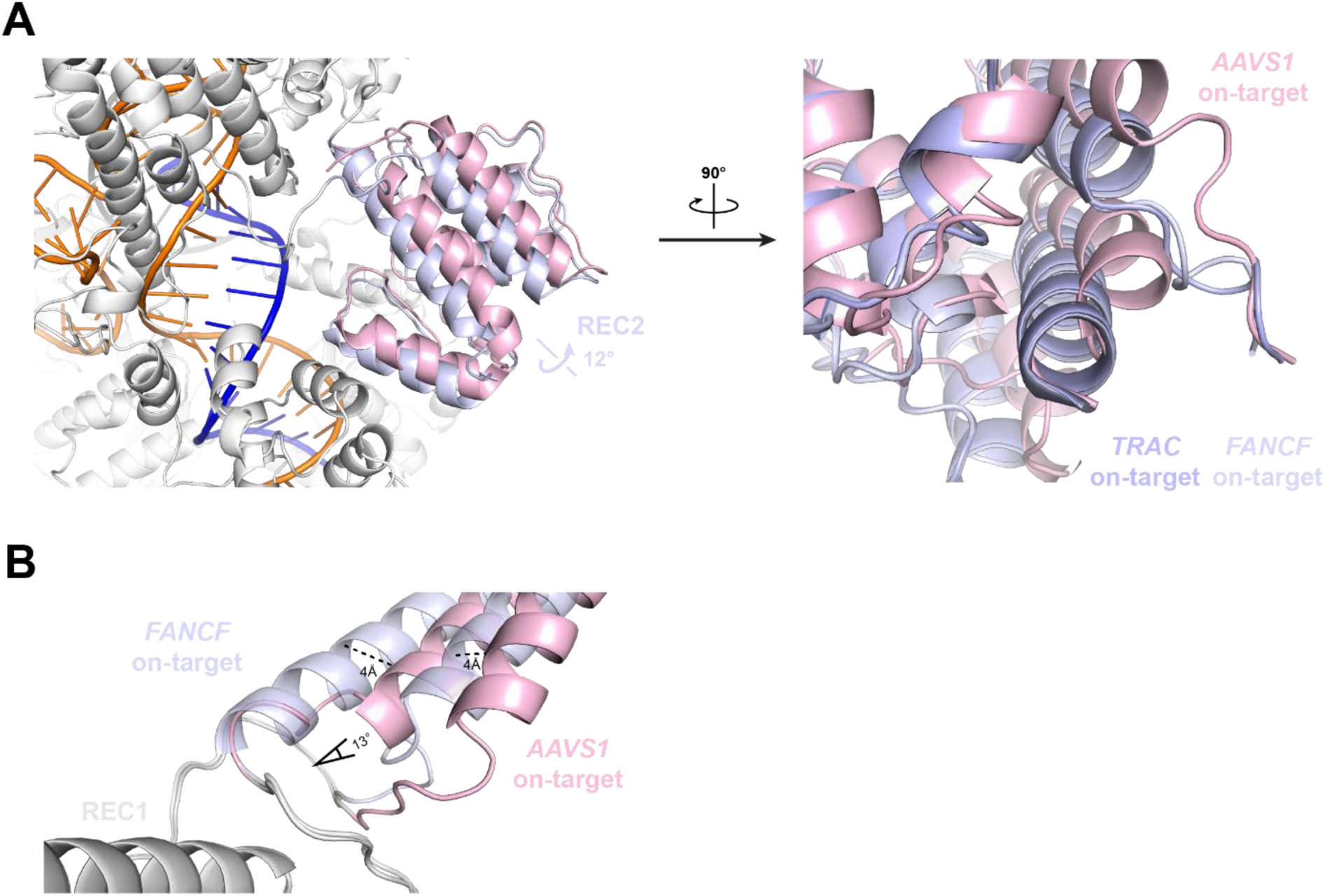
Alternative REC2 conformation in *AAVS1* on-target complex. (**A**) Overlay of REC2 domain conformations in the *AAVS1* (pink), *FANCF* (purple) and *TRAC* (light blue) on-target complexes (**B**) Close-up view of helix REC2 helix spanning Cas9 residues 174-180. Linear and angular displacements of the helix in the *AAVS1* on-target complex relative to the *FANCF* and *TRAC* on-target complexes are indicated.

**Figure S5.**
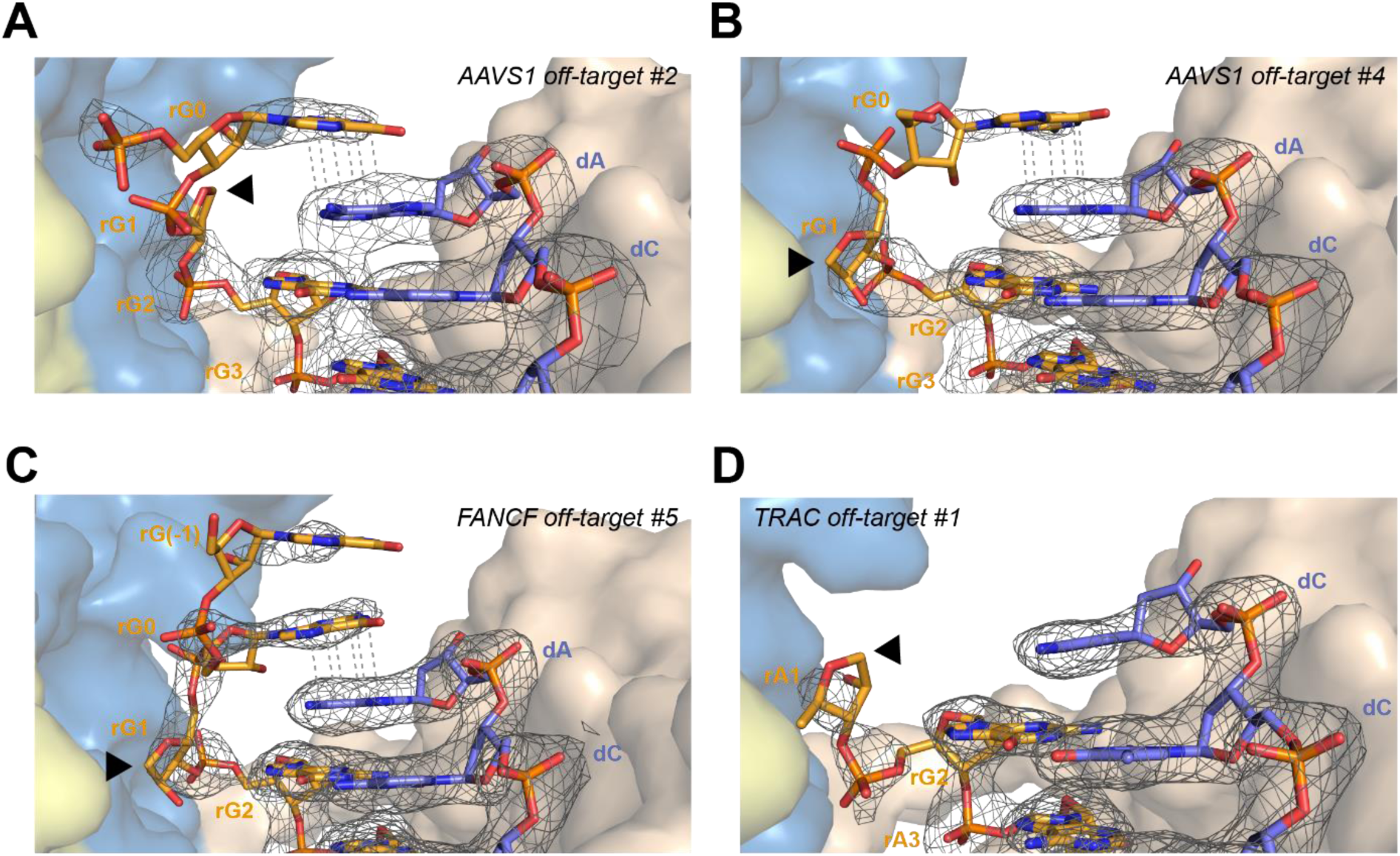
PAM-distal mismatches result in unpairing and disordering of guide RNA nucleobase in position 1. Close-up views of the PAM-distal end of the guide RNA-TS heteroduplex in (**A**) *AAVS1* off-target #2, (**B**) *AAVS1* off-target #4, (**C**) *FANCF* off-target #5 and (**D**) *TRAC* off-target #1 complexes. Arrowheads indicate nucleotides with disordered bases. Refined 2m*F*_o_−D*F*_c_ electron density maps of the heteroduplexes are rendered as a grey mesh and contoured at 1.2σ for (A) and 1.0σ for (B)-(D).

**Figure S6.**
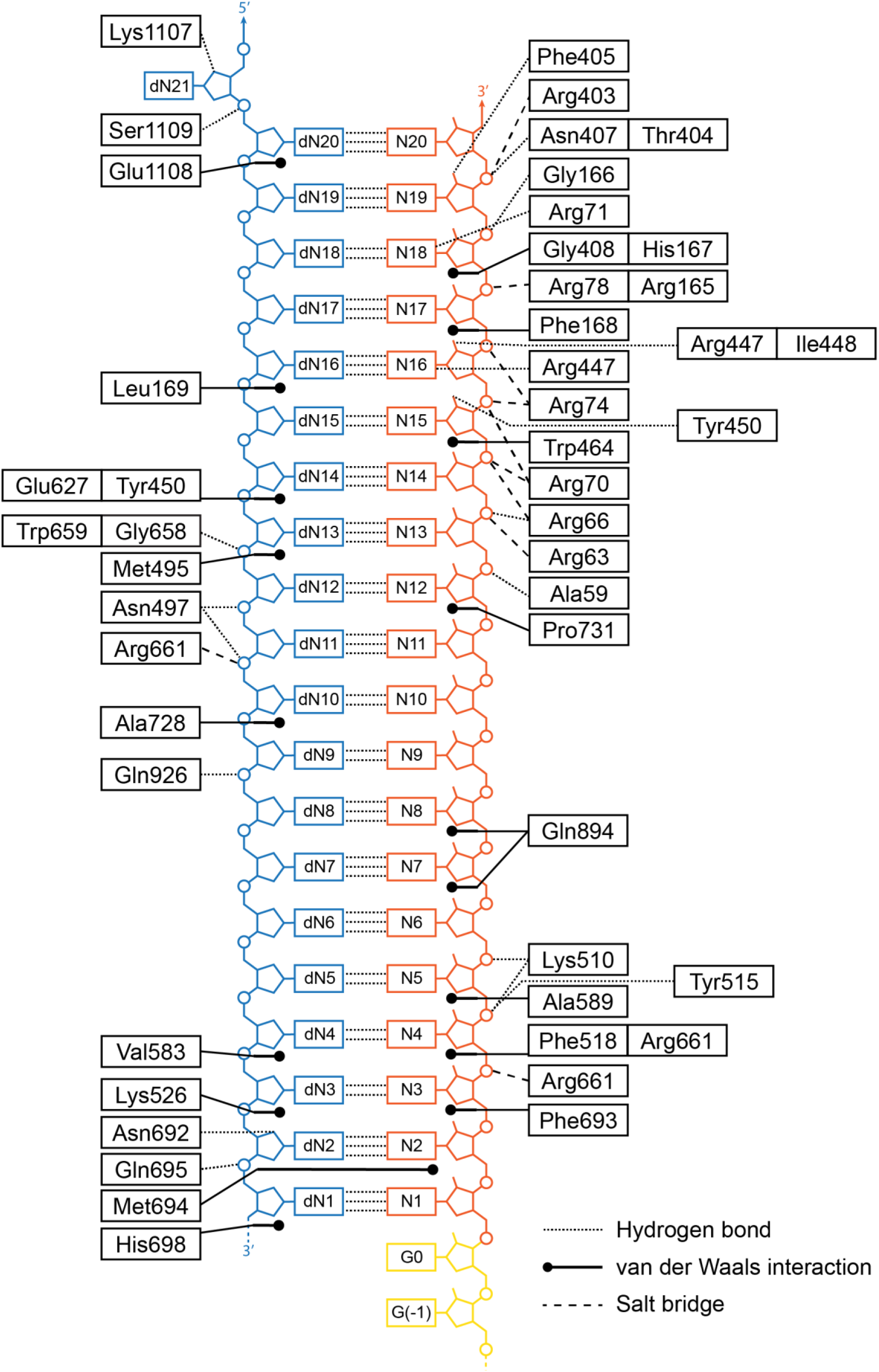
Cas9-nucleic acid interactions in on-target complexes. Schematic diagram depicting Cas9 residues interacting with the guide RNA-target DNA heteroduplex. Dotted lines represent hydrogen bonding interactions; dashed lines represent salt bridges; solid lines represent stacking/hydrophobic interactions. Target strand is coloured in blue, guide RNA in orange. Phosphates are represented by circles, ribose moieties by pentagons, and nucleobases by rectangles.

**Figure S7.**
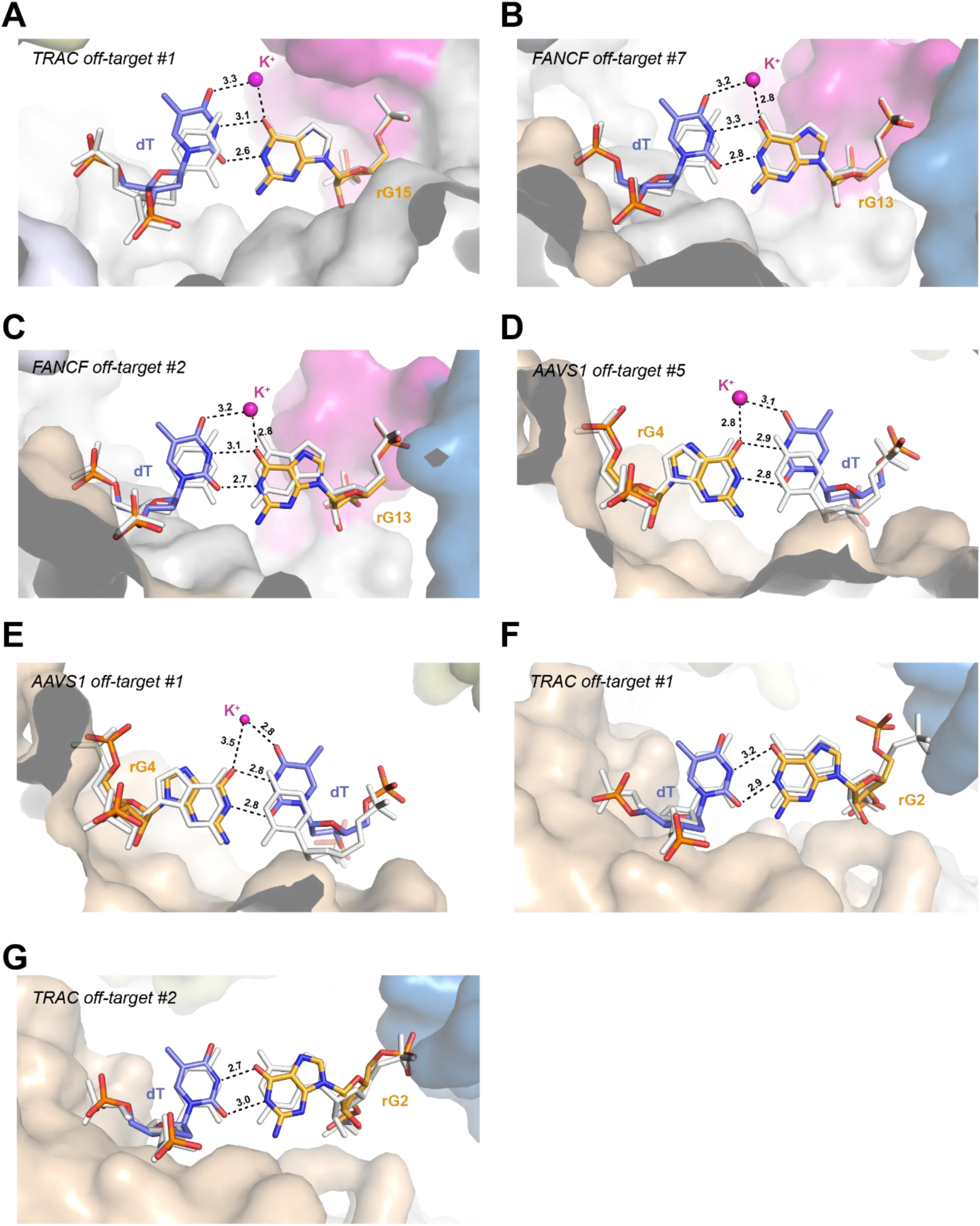
Wobble base pairing of rG-dT mismatches. Close-up views of rG-dT mismatches at (**A**) heteroduplex position 15 in the *TRAC* off-target #1 complex, (**B**) position 13 in *FANCF* off-target #7 complex, (**C**) position 13 in *FANCF* off-target #2 complex, (**D**) position 4 in *AAVS1* off-target #5 complex, (**E**) position 4 in *AAVS1* off-target #1 complex, (**F**) position 2 in *TRAC* off-target #1 complex and (**G**) position 2 in *TRAC* off-target #2 complex. Corresponding on-target Watson-Crick base pairs are shown in white. Monovalent ions, modeled as K^+^, are depicted as purple spheres. In (A)-(E), the dT base is displaced into the major groove and forms a canonical wobble base pair with the rG base. In (F)-(G), the the rG base instead shifts towards the minor groove to facilitate wobble pairing.

**Figure S8.**
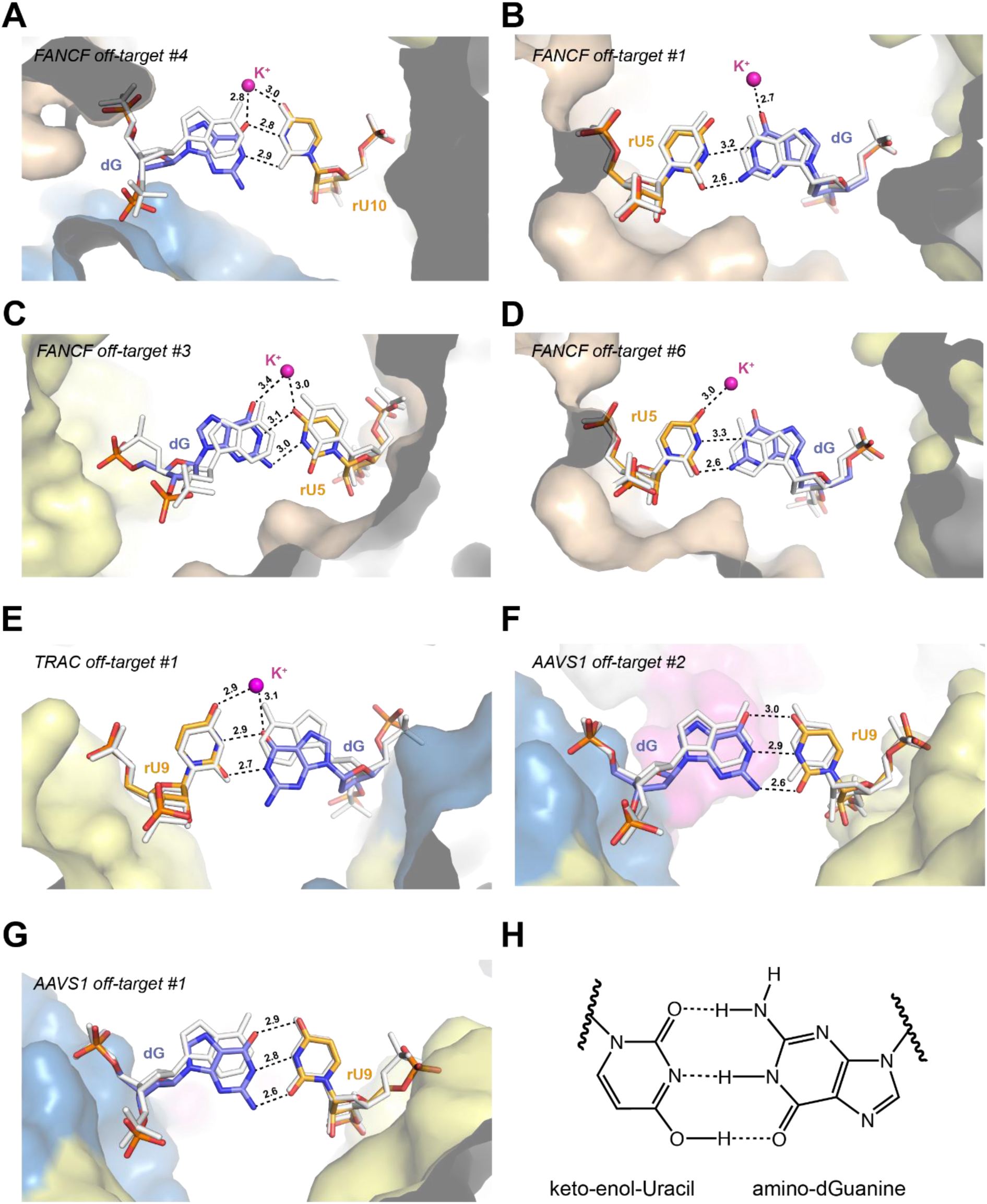
rU-dG wobble base pairs adopt duplex position-specific conformations. Close-up views of of rU-dG mismatches at (**A**) heteroduplex position 10 in *FANCF* off-target #4 complex, (**B**) position 5 in *FANCF* off-target #1 complex, (**C**) position 5 in *FANCF* off-target #3 complex, **(D**) position 5 in *FANCF* off-target #6 complex, (**E**) *TRAC* off-target #1 complex, and (**F**) *AAVS1* off-target #2, and (**G**) *AAVS1* off-target #1 complex. Corresponding on-target Watson-Crick base pairs are shown in white. Bound potassium ions are depicted as purple spheres. In (A), rU-dG wobble base pairing is achieved by minor groove displacement of the guanine base. In (B)-(D), the rU-dG mismatches adopt atypical conformations. In (E), the guanine base is shifted into the minor groove to form a wobble base pair, whereas at the identical heteroduplex position in (F) and (G), the rU-dG base pairs do not engage in wobble pairing, instead adopting alternative tautomeric forms. **(H)** Schematic depicting hydrogen bonding interactions between rU and dG bases in (F) and (G).

**Figure S9.**
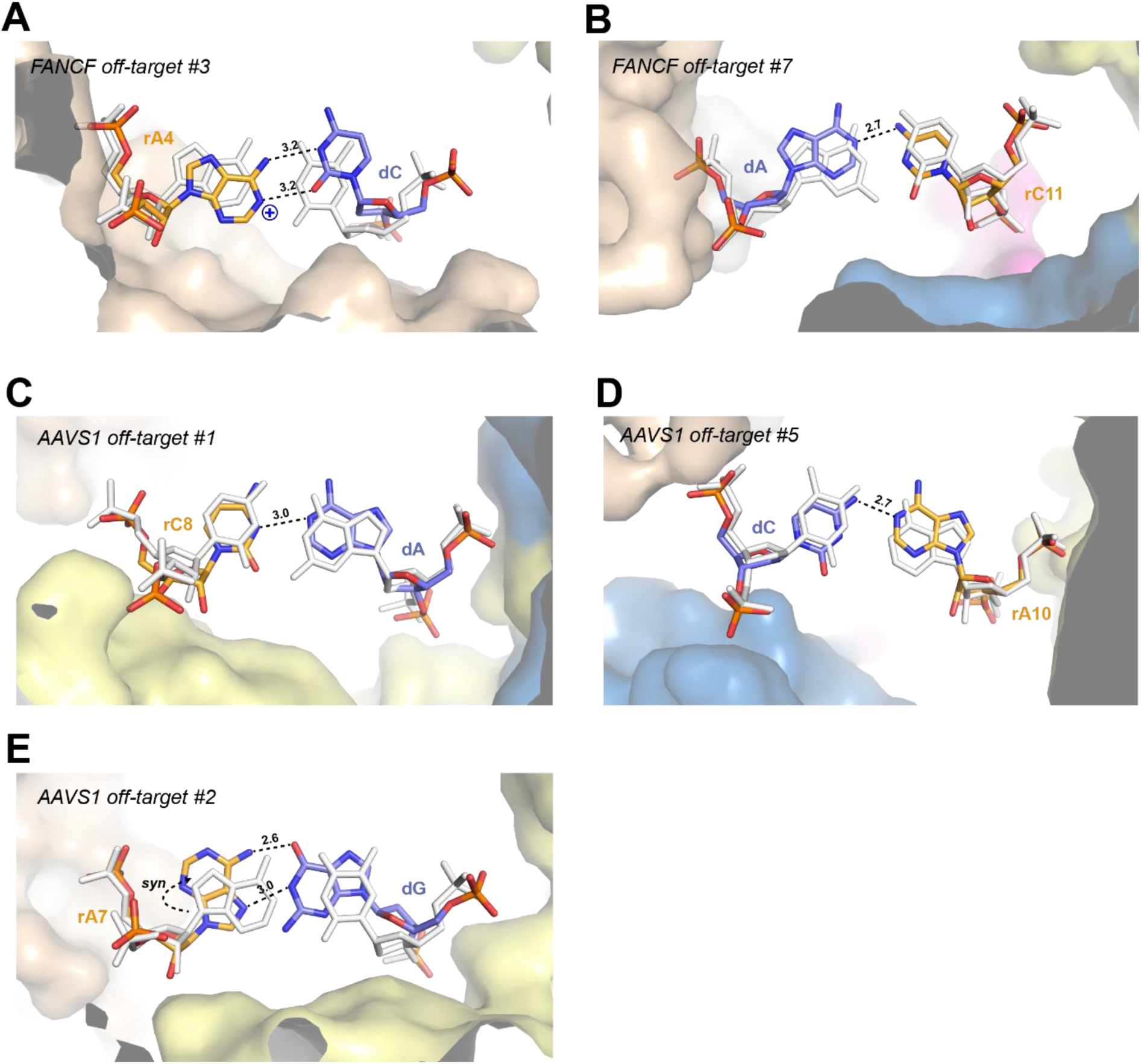
Non-canonical pyrimidine-purine base pairs within Cas9 off-target complexes. (**A**) Close-up view of rA-dC wobble base pairing at position 4 in *FANCF* off-target #3 complex. The base pair geometry is consistent with base protonation or tautomerism to enable productive hydrogen bonding between the bases. (**B**) Close-up view of rC-dA mismatch at position 11 of *FANCF* off-target #7 complex, facilitated by base tilting at positions 11 and 12. (**C**) Close-up view of partially paired rC-dA mismatch at position 8 in *AAVS1* off-target #1 complex. (**D**) Close-up view of rA-dC mismatch at position 10 in *AAVS1* off-target #5 complex. (**E**) Close-up view of Hoogsteen-edge rA-dG base pair at position 7 in *AAVS1* off-target #2 complex. Corresponding on-target Watson-Crick base pairs are shown in white.

**Figure S10.**
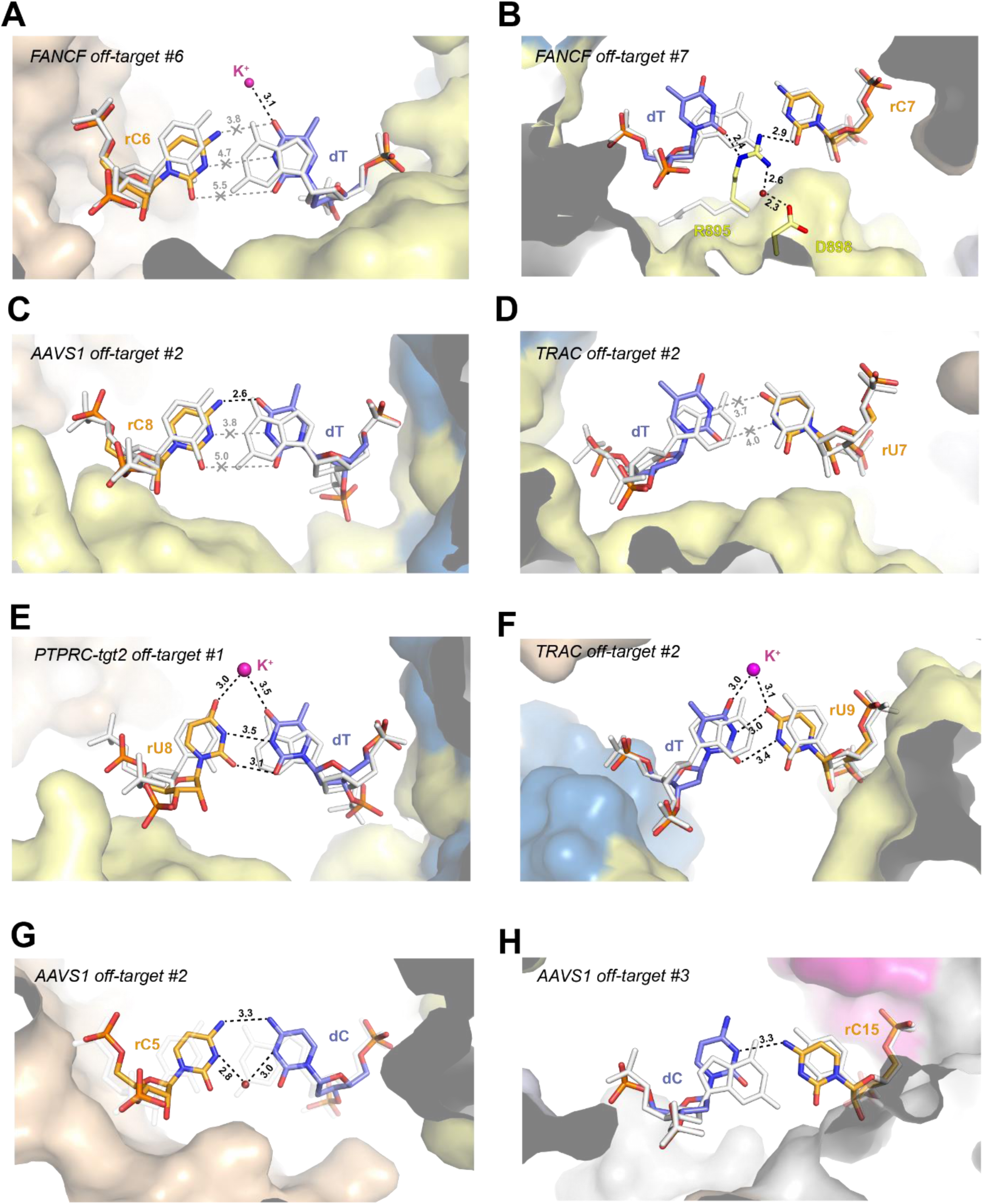
Pyrimidine-pyrimidine mismatches within off-target complexes. (**A**) Close-up view of rC-dT mismatch at position 6 in *FANCF* off-target #6 complex. (**B**) Close-up view of rC-dT mismatch at position 7 in *FANCF* off-target #7 complex, bridged by Arg895. The arginine sidechain in the corresponding on-target complex is shown in white. (**C**) Close-up view of rC-dT base pairing at position 8 of *AAVS1* off-target #2. (**D**) Close-up view of rU-dT mismatch at position 7 in *TRAC* off-target #2 complex. (**E**) Close-up view of rU-dT pairing at position 9 in *PTPRC*-tgt2 off-target #1 complex, facilitated by base propeller twisting. (**F**) Close-up view of rU-dT pairing at position 8 in *TRAC* off-target #2 complex, enabled by backbone shift of the RNA strand. (**G**) Close-up view of partially paired rC-dC mismatch at position 5 in *AAVS1* off-target #2 complex, bridged by a water molecule. (**H**) Close-up view of rC-dC mismatch at position 15 in *AAVS1* off-target #3 complex. Corresponding on-target Watson-Crick base pairs are shown in white (for *PTPRC-tgt2*, the *FANCF* on-target structure was used and bases were mutated *in silico*).

**Figure S11.**
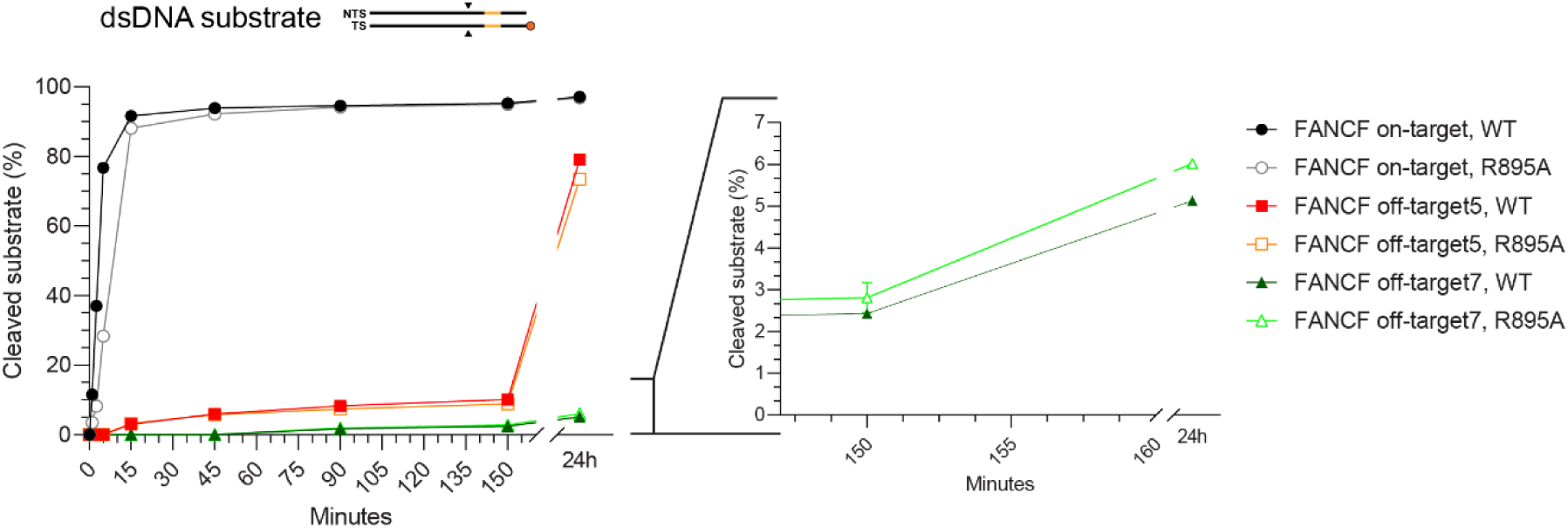
Cas9 R895A mutation of Cas9 does not affect FANCF off-target #7 cleavage or binding. (**A**) Kinetic analysis of *FANCF* on- and off-target substrate DNA cleavage by wild-type and R895A Cas9 proteins.

**Figure S12.**
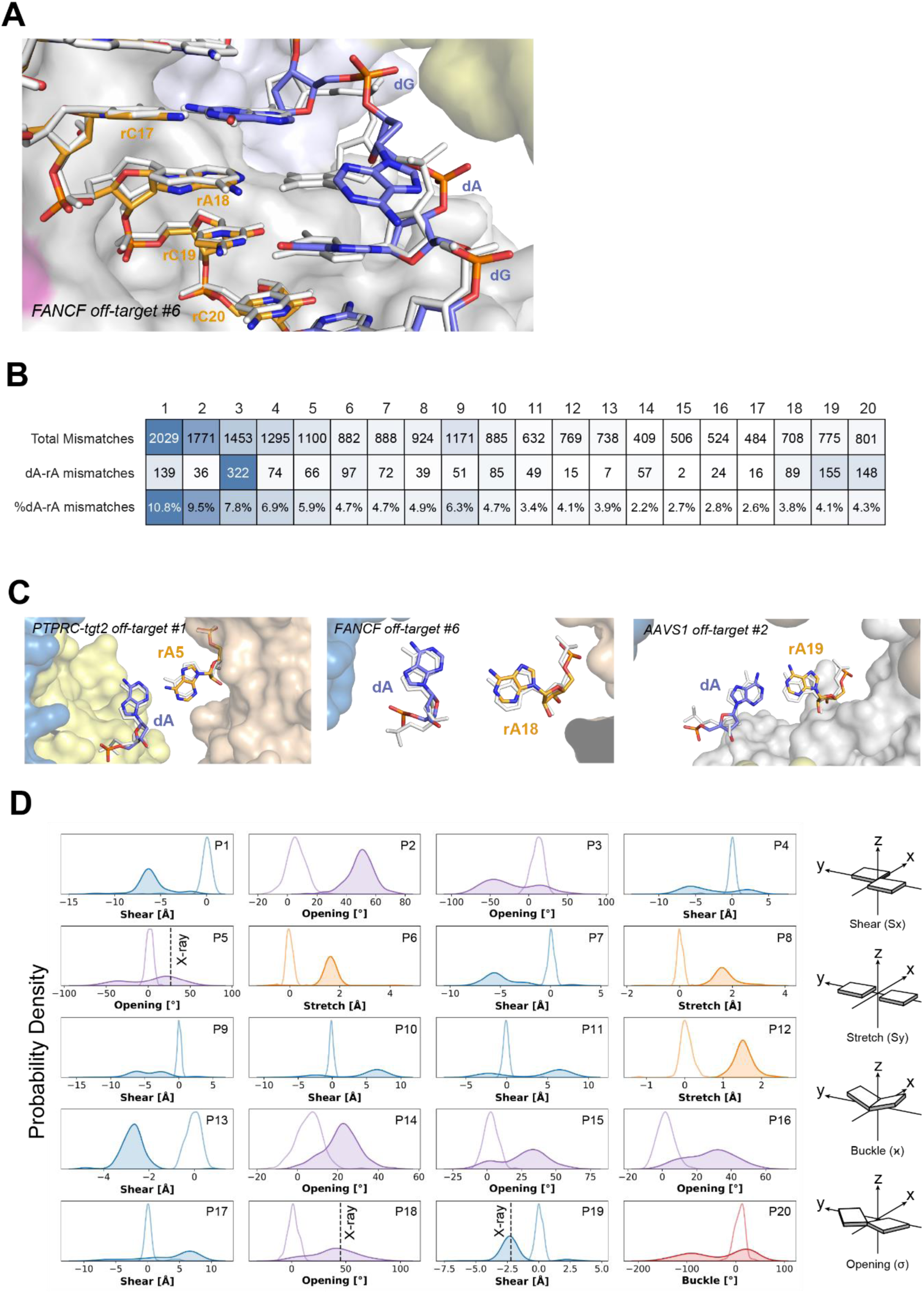
Tolerance of rA-dA mismatches within the heteroduplex. (**A**) Close-up view of rA-dA mismatch at position 18 in *FANCF* off-target #6 complex, overlaid with the *FANCF* on-target structure (white). (**B**) Number of rA-dA off-target mismatches per heteroduplex position recovered in the SITE-Seq assay for all analysed genomic targets. Percentages indicate frequency of rA-dA mismatches recovered in the particular position as a fraction of total number of rA-dA mismatches. (**C**) Close-up view of MD-simulated rA-dA mismatches (colored), overlaid with crystallographic ground truth (white) at positions 5 (PTPRC-tgt2 off-target #1), 18 (FANCF off-target #6), and 19 (AAVS1 off-target #2). (**D**) Probability density plots of the most representative geometrical base pair parameters (viz., shear in blue, opening in violet, stretch in orange and buckle in red), describing the mode of rA-dA mismatch tolerance at different positions (1 – 20) of the heteroduplex, as computed via MD simulations for the systems with (shaded) and without mismatches (unshaded). Vertical line in positions 5, 18 and 19 indicates base pair parameter value of the respective X-ray structure. For each system, data are reported considering an aggregate sampling of ~1.5 μs.

**Figure S13.**
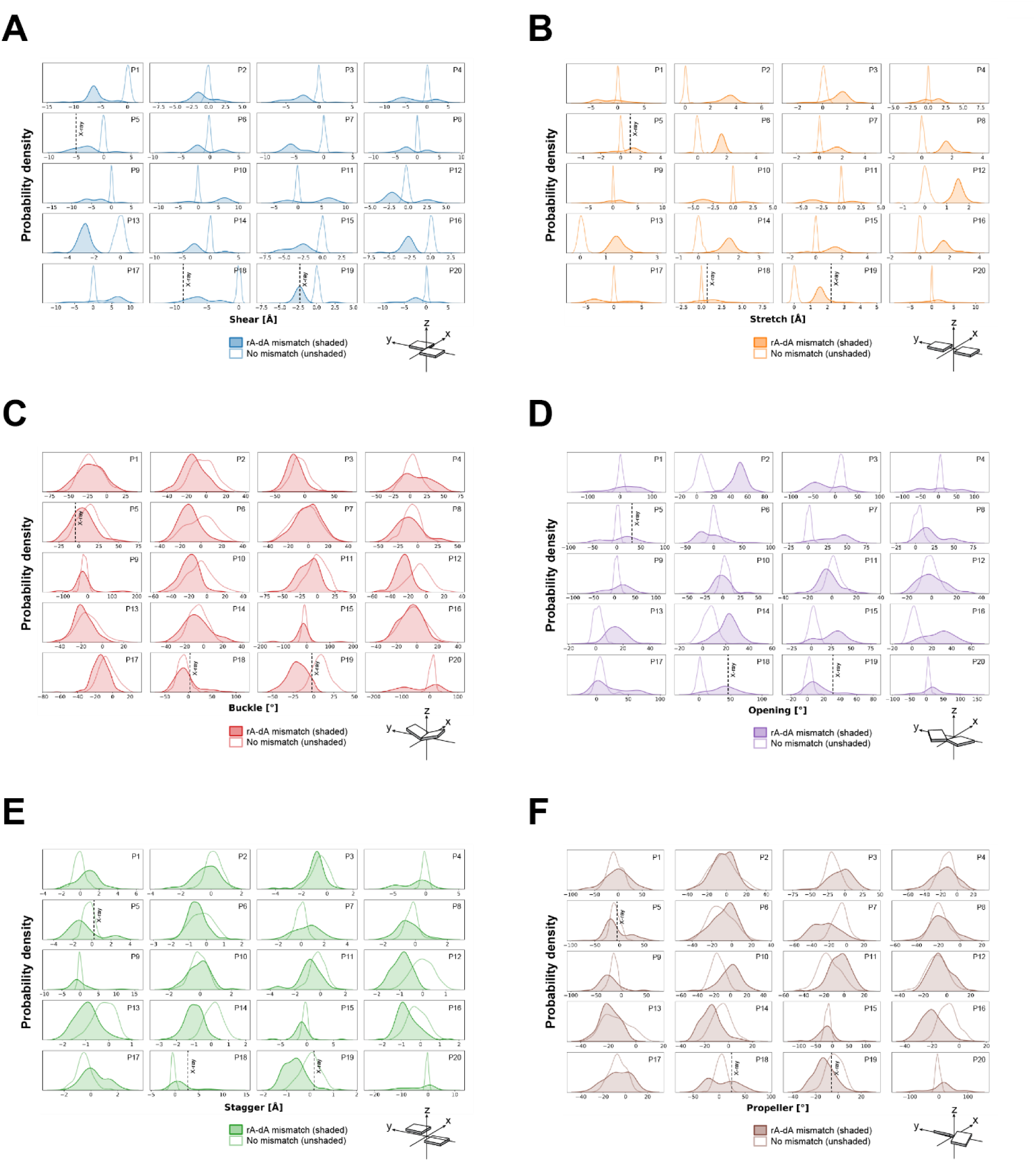
Base pair parameters of MD simulated rA-dA mismatches. Probability density plots for (**A**) shear, (**B**) stretch, (**C**) buckle, (**D**) opening, (**E**) stagger, and (**F**) propeller-twist base pair parameter for the rA-dA mismatch at different heteroduplex, as computed from molecular dynamics simulations for the systems with (shaded) and without mismatches (unshaded). For each system, data are reported considering an aggregate sampling of ~1.5 μs.

**Figure S14.**
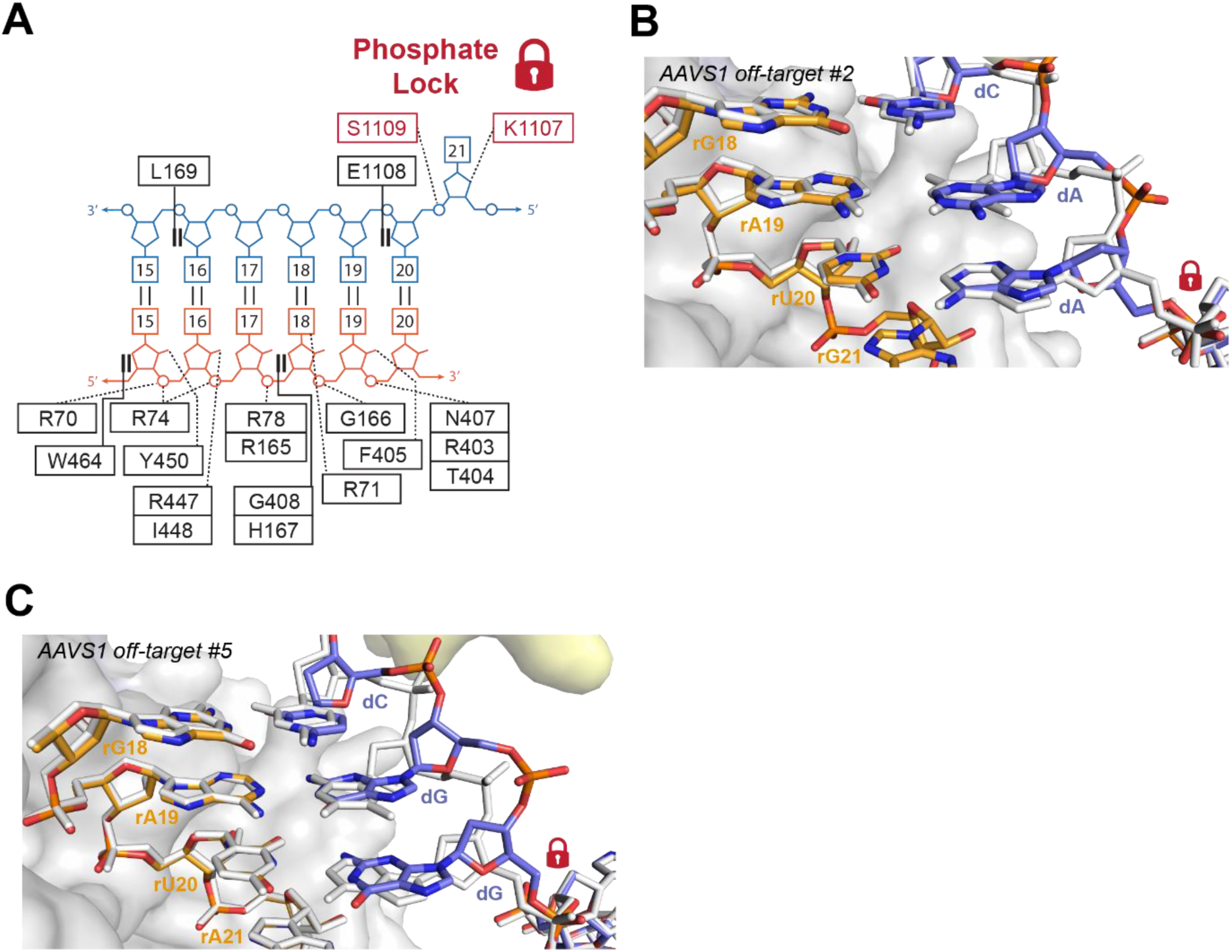
Lack of protein contacts with the target DNA strand in the seed region facilitates backbone distortions. (**A**) Schematic overview of Cas9 interactions within the PAM-proximal seed region of the guide RNA-TS DNA heteroduplex. (**B**) Close-up view of the seed region in *AAVS1* off-target #2 complex, overlaid with the AAVS1 on-target heteroduplex (white), showing structural distortion of the TS due to rA-dA mismatch at seed position 19. (**C**) Close-up view of the seed region in AAVS1 off-target #5 complex, overlaid with the AAVS1 on-target heteroduplex (white), showing structural distortion due to rA-dG and rU-dG mismatches at positions 19 and 20, respectively. Red lock icon indicates position of the phosphate lock residue in (B) and (C).

**Figure S15.**
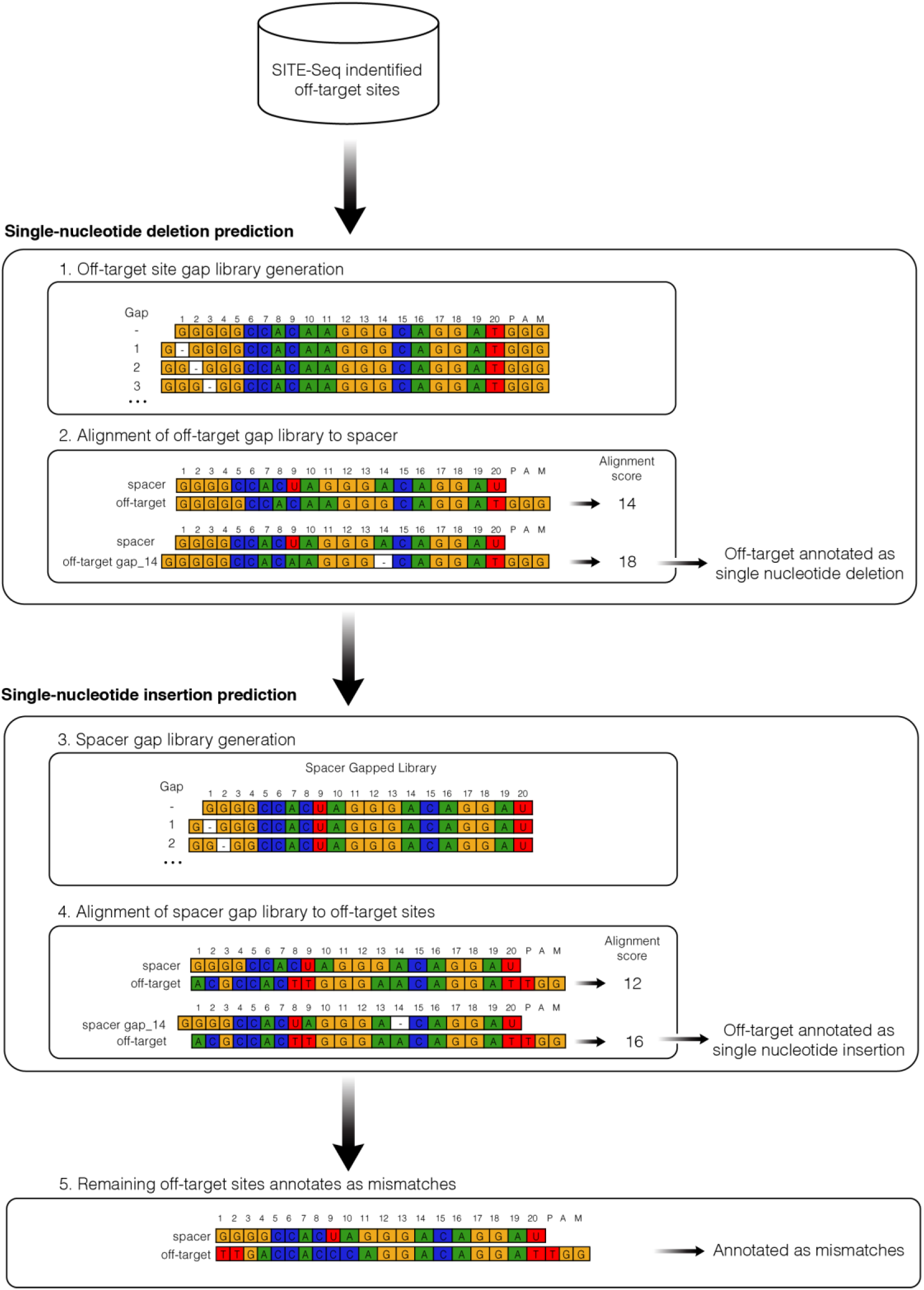
Schematic representation of mismatch, insertion, and deletion classification algorithm for the SITE-Seq off-target analysis. Schematic represents classification algorithm of off-target sites with putative insertions and deletions.

**Figure S16.**
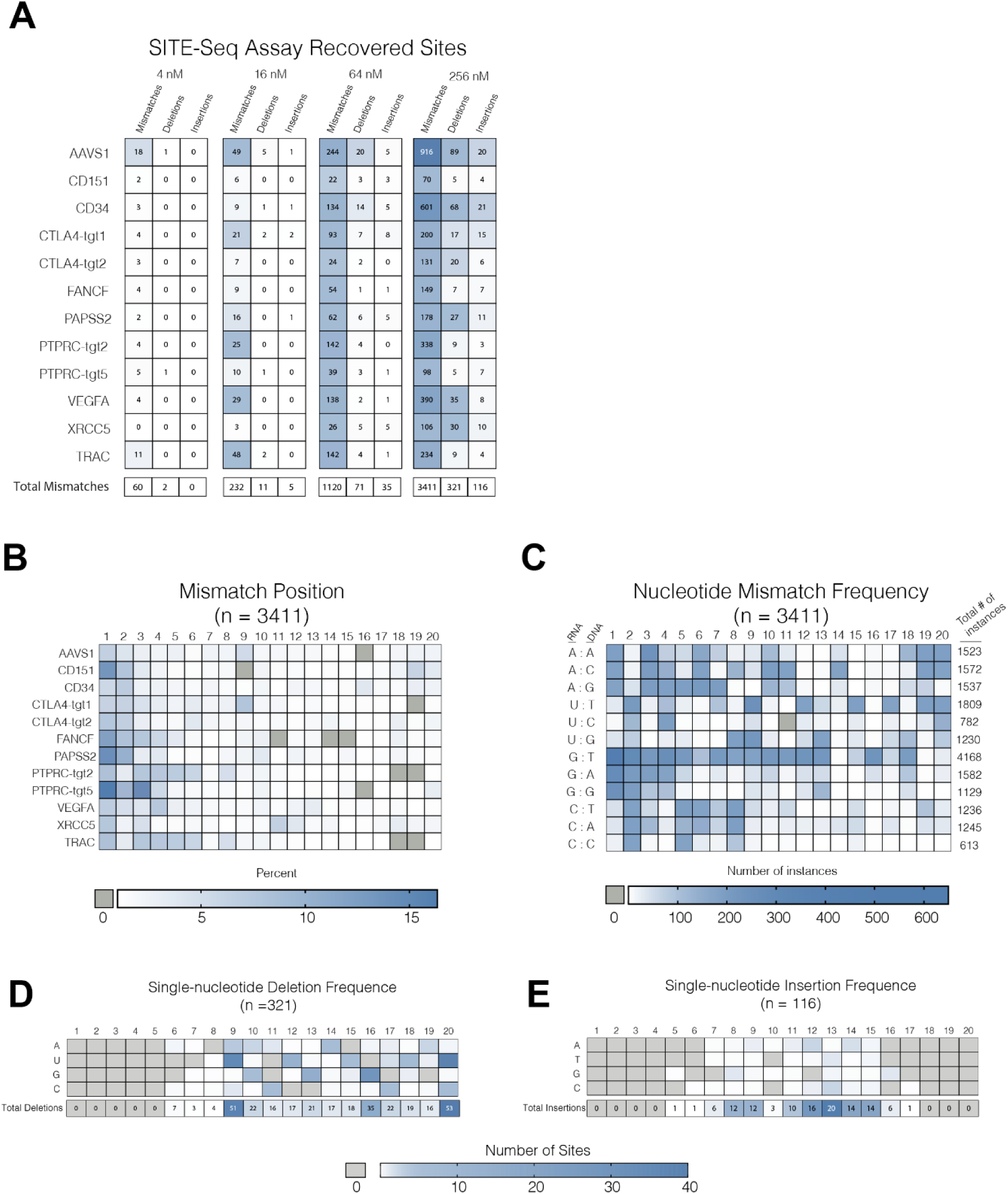
Positional mapping of nucleotide insertions and deletions in SITE-Seq off-target analysis. (**A**) Number of recovered off-target sites per genomic target as a function of RNP concentration classified as containing either only mismatches, single-nucleotide deletions, or single-nucleotide insertions. (**B**) Frequency of positional mismatch occurrence per genomic target for mismatched off-targets. (**C**) Frequency of nucleotide mismatches within the heteroduplex for all off-target sites (n=3411 sites for both (B) and (C)). (**D**) Frequency of single-nucleotide deletions for all off-target sites. (n=321 sites). (**E**) Frequency of single-nucleotide insertions for all off-target sites. (n=116 sites).

**Figure S17.**
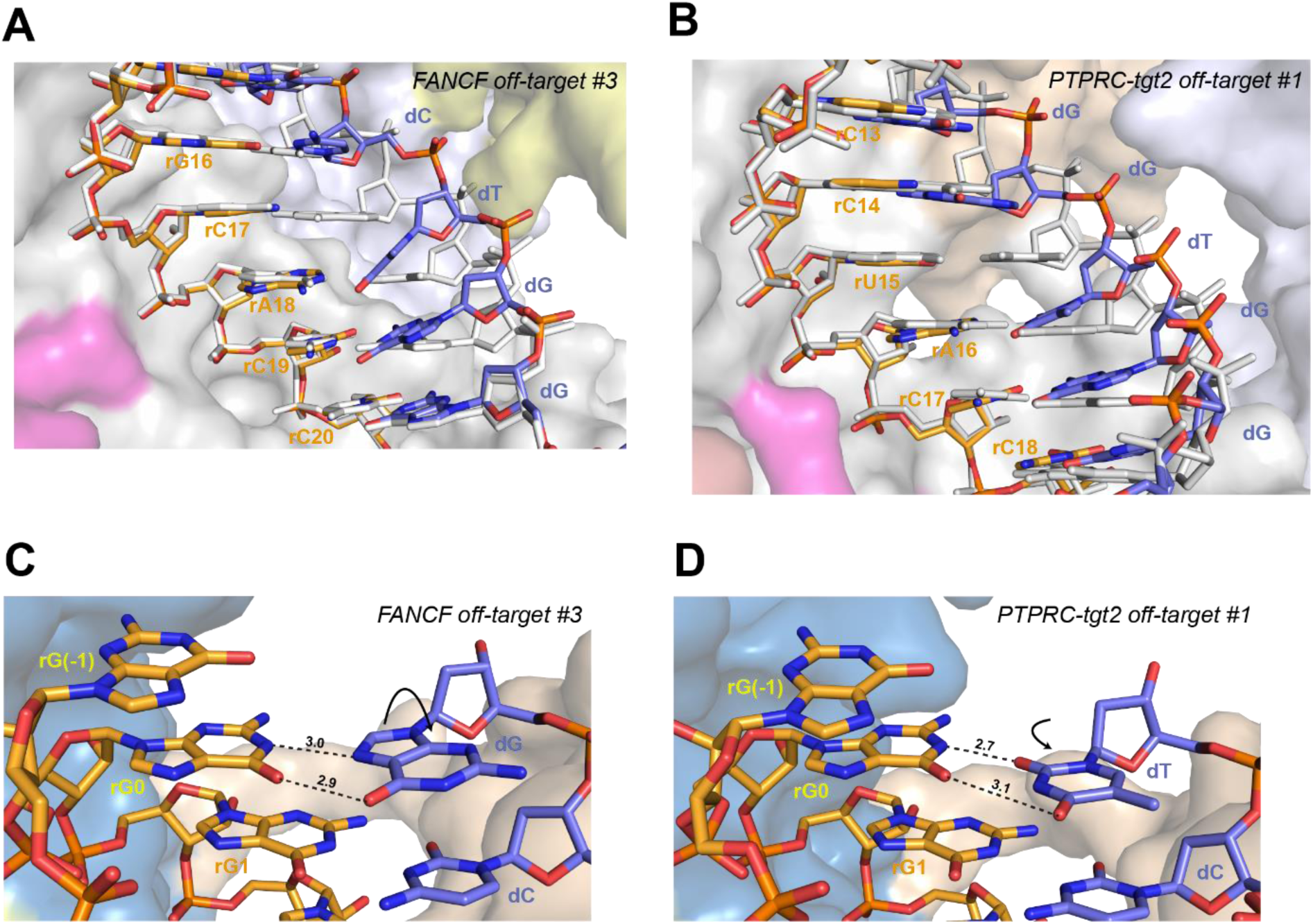
Recognition of off-target sites containing deletions in the seed region. (**A**) Close-up view of base skipping within the seed region of the guide RNA-off-target DNA heteroduplex in *FANCF* off-target #3 complex, overlaid with the on-target heteroduplex (white). (**B**) Close-up view of base skipping within the seed region of the guide RNA-off-target DNA heteroduplex in *PTPRC-tgt2* off-target #1 complex, overlaid with *FANCF* on-target heteroduplex (white). (**C**) Close-up view of non-canonical base pairs at the 5’-terminus of the guide RNA in *FANCF* off-target #3 complex involving guanosine nucleotides introduced during *in vitro* transcription of the guide RNA. (**D**) Close-up view of non-canonical base pairs at the 5’-terminus of the guide RNA in *PTPRC-tgt2* off-target #1 complex involving guanosine nucleotides introduced during *in vitro* transcription of the guide RNA.

**Figure S18.**
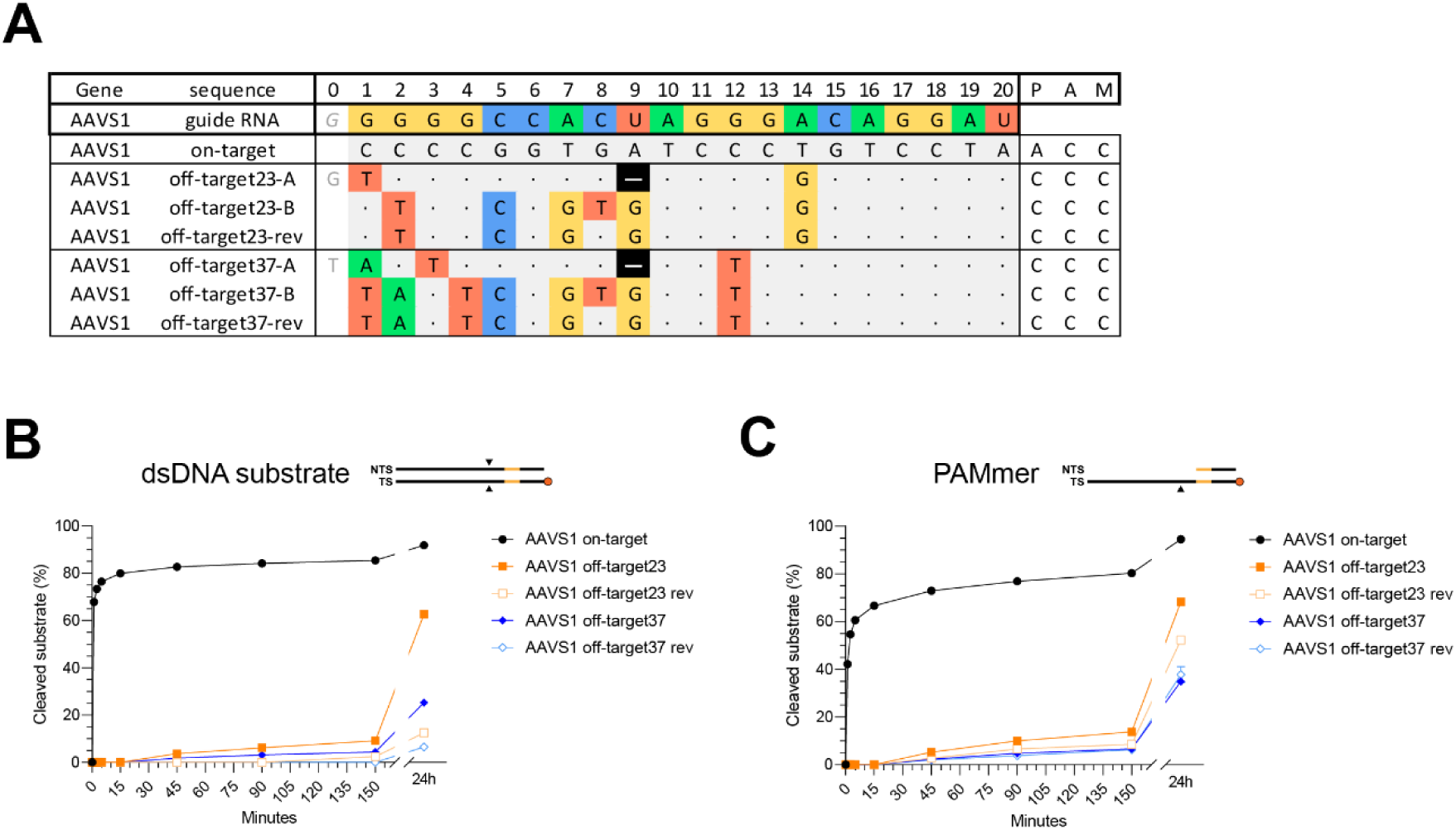
Mismatch reversal reduces cleavage efficiency of off-targets containing multiple consecutive mismatches. (**A**) Schematic depiction of possible mismatch tolerance mechanisms through PAM-distal deletions (denoted-A) or the formation of multiple mismatches (denoted-B) in *AAVS1* off-targets. Off-targets denoted with “-rev” indicate the reversal of a single mismatch in the consecutive mismatch region back to the corresponding canonical base pair. (**B**) *In vitro* cleavage kinetics of fully double stranded on- and off-target DNA substrates for off-targets with putative multiple consecutive mismatches in the PAM-distal region. Black triangles in the substrate schematic (top) indicate position of cleavage sites. Each data point represents a mean of four independent replicates. Error bars represent standard deviation for each time point. (**C**) *In vitro* cleavage kinetics of corresponding partially single stranded (PAMmer) on- and off-target DNA substrates

**Figure S19.**
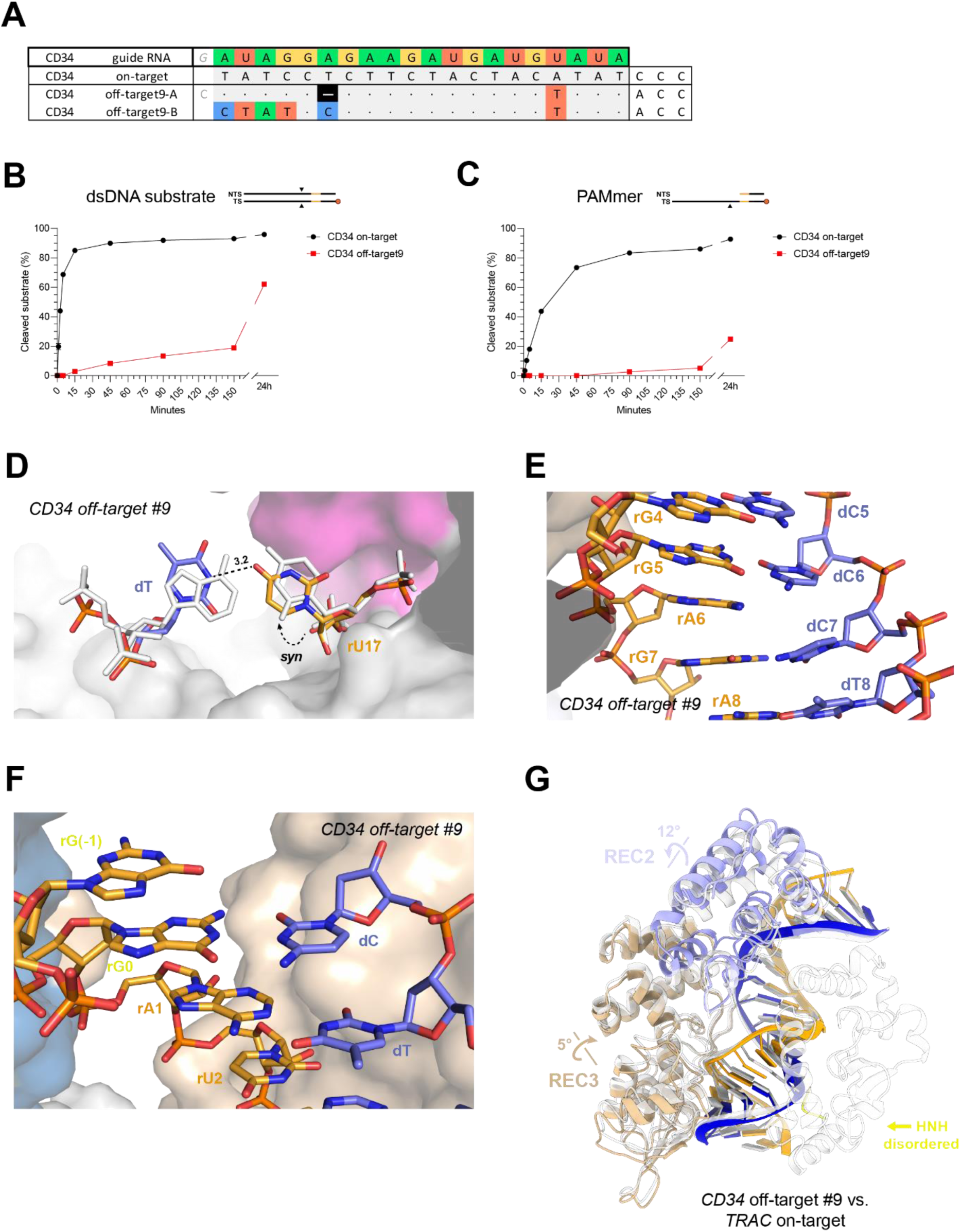
Base skipping in off-target substrate containing PAM-distal deletion. (**A**) Schematic depiction of a PAM-distal single-nucleotide deletion in the *CD34* off-target #9 sequence. (**B**) *In vitro* cleavage kinetics of fully double stranded *CD34* on-target and off-target #9 substrates. Black triangles in the substrate schematic (top) indicate position of cleavage sites. Each data point represents a mean of four independent replicates. Error bars represent standard deviation for each time point. (**C**) In vitro cleavage kinetics of partially single stranded (PAMmer) *CD34* on-target and off-target #9 substrates. (**D**) Close-up view of rU-dT mismatch at position 17 in *CD34* off-target #9 complex, where the rU17 nucleotide undergoes *anti-syn* isomerisation. Corresponding on-target Watson-Crick base pairs are shown in white (the *TRAC* on-target structure was used and bases were mutated *in silico*). (**E**) Zoomed-in view of the PAM-distal base skip at duplex position 6 in the *CD34* off-target #9 complex. (**F**) Close-up view of rG-dC Watson-Crick base pair at the 5’-terminus of the guide RNA in *CD34* off-target #9 complex, involving guanosine nucleotides introduced during *in vitro* transcription of the guide RNA. (**G**) Structural overlay of the *CD34* off-target #9 complex with the *TRAC* on-target complex. The REC1, RuvC, and PAM interaction domains have been omitted for clarity in all panels. The *CD34* off-target #9 complex domains are colored according to **Figure 1A**. The on-target complex is colored white.

**Figure S20.**
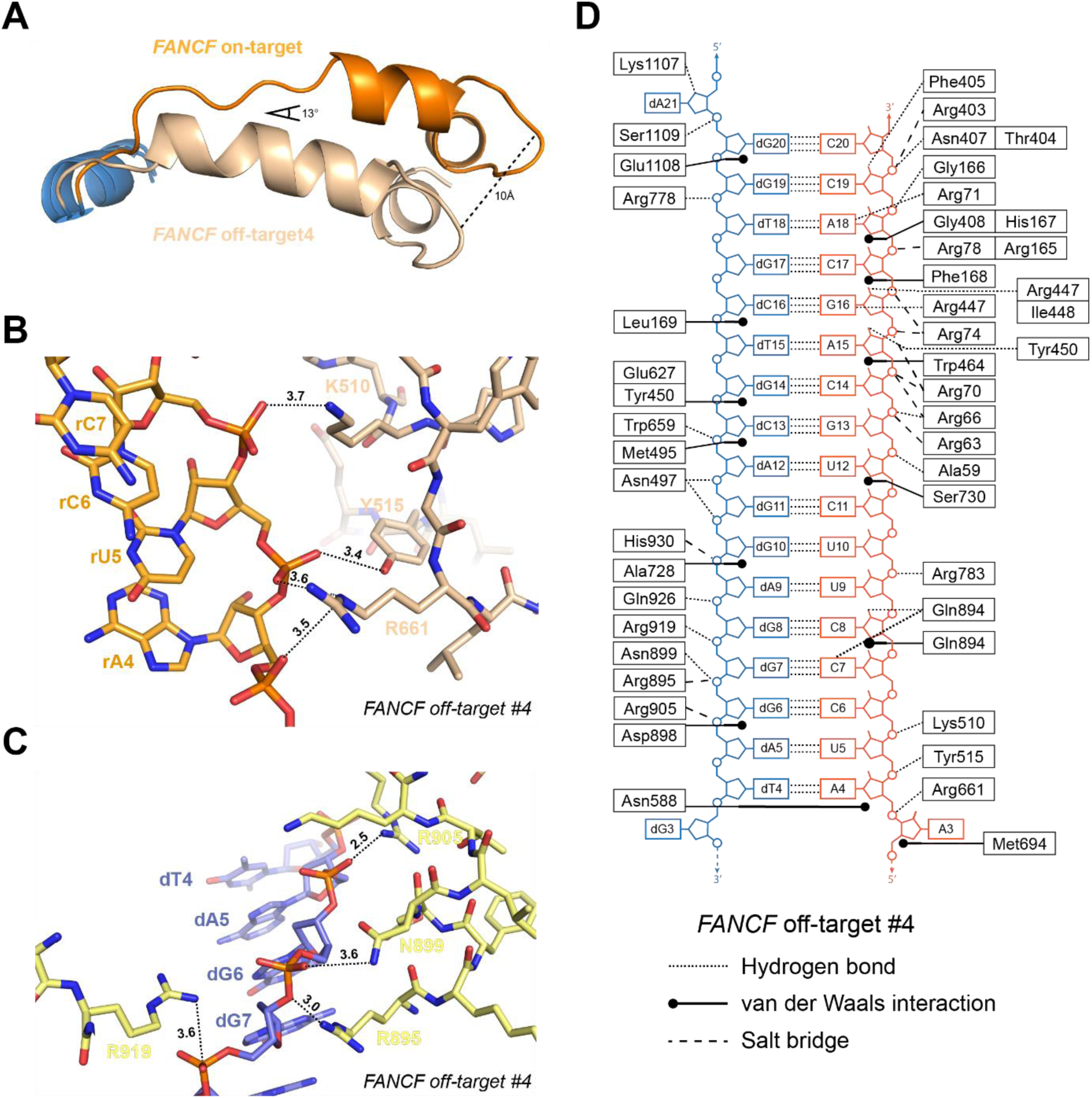
Altered heteroduplex interactions in *FANCF* off-target #4 complex. (**A**) Overlay of REC3 domain helix 703-712 in *FANCF* off-target #4 complex (wheat) with *FANCF* on-target complex (orange). (**B**) Close-up view of REC3 domain interactions with the guide RNA strand in *FANCF* off-target #4 complex. (**C**) Close-up view of TS DNA interactions established by HNH domain in *FANCF* off-target #4 complex. (**D**) Schematic diagram depicting Cas9 residues interacting with the guide RNA-off-target DNA heteroduplex in *FANCF* off-target #4 complex. Dotted lines represent hydrogen bonding interactions, dashed lines represent salt bridges, solid lines represent stacking/hydrophobic interactions. Target strand is coloured blue, guide RNA orange. Phosphates are represented by circles, ribose moieties by pentagons, and nucleobases by rectangles.

**Figure S21.**
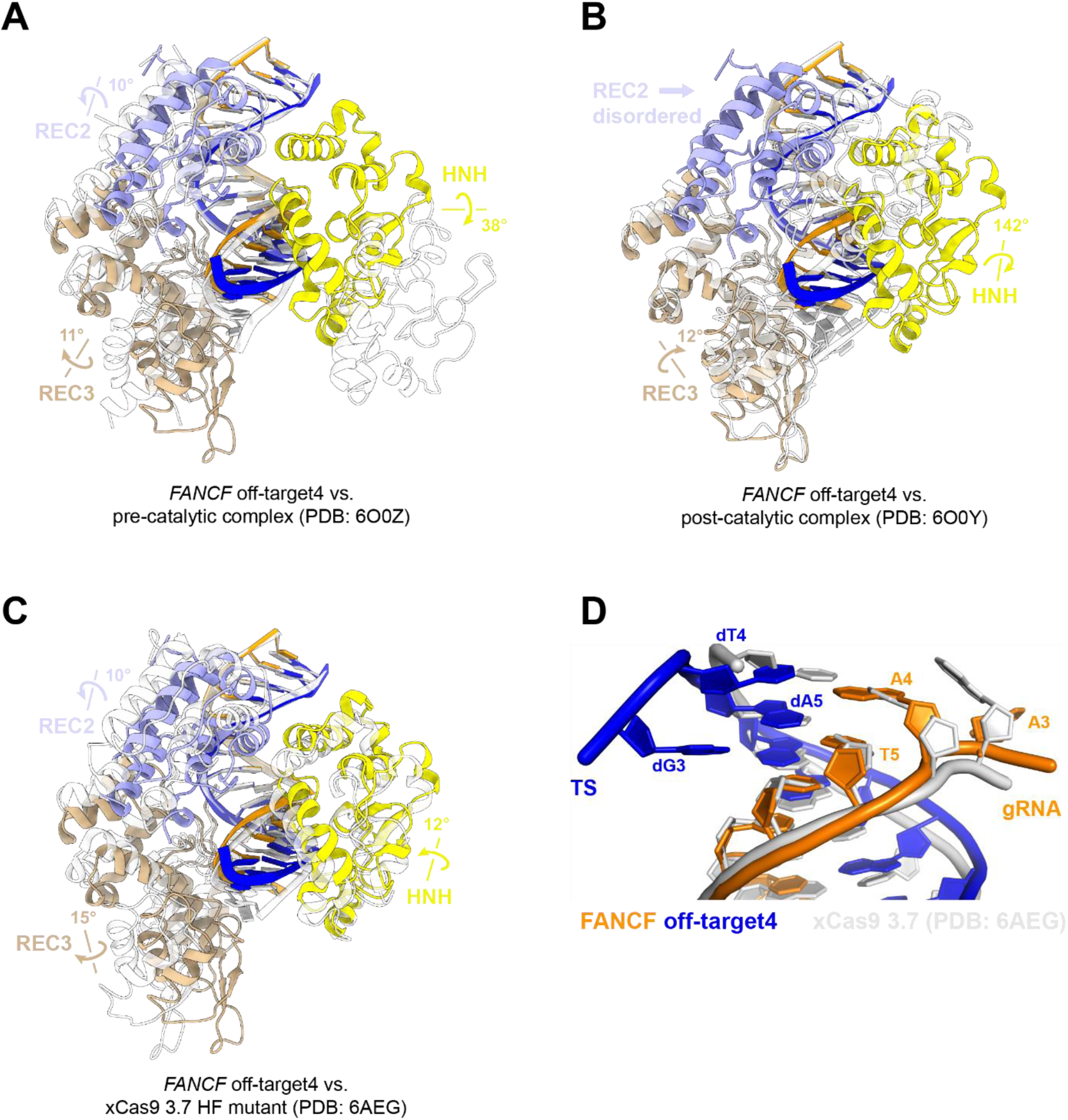
Conformational rearrangements of REC2/3 AND HNH domains in *FANCF* off-target #4 complex. (**A**) Structural overlay of the *FANCF* off-target #4 complex with cryo-EM structure of a pre-catalytic (State I) Cas9 complex (PDB: 6O0Z). (**B**) Structural overlay of the *FANCF* off-target #4 complex with the cryo-EM structure of a post-catalytic (State II) Cas9 complex (PDB: 6O0Y). (**C**) Structural overlay of the *FANCF* off-target #4 complex with the crystallographic structure of the high-fidelity xCas9 3.7 variant (PDB: 6AEG). The REC1, RuvC, and PAM interaction domains have been omitted for clarity in all panels, as no significant structural changes were observed in these domains. The *FANCF* off-target #4 complex domains are colored according to **Figure 1A**. The overlaid structures are coloured white. (**D**) Overlay of the PAM-distal heteroduplex region in *FANCF* off-target #4 and xCas9 3.7 on-target complexes. Target strand is coloured in blue; guide RNA is coloured orange.

**Table S1. SITE-Seq assay analysis of Cas9 off-target sites for 12 selected genomic targets. Related to Figure S1**

Columns indicate the recovered off-target sequence; motif location; number of substitutions in recovered target sequence compared to the on-target (substitutions); strand designation of PAM; the lowest recovery concentration of each target; and whether the off-target is predicted to contain inserts or deletions based. Off-target sites recovered at lower concentrations were also recovered at higher concentrations (*e.g.,* all 4 nM sites were also recovered at 16 nM, 64 nM, and 256 nM).

**Table S2. List of recovered off-target sites by SITE-Seq. Related to Figures S1 and S19**

Off-target alignments classified by genomic target and by the presence of insertions, deletions or purely mismatched targets. Indexes correspond to off-target sequence numbering in Table S1.

**Table S3. Cellular editing data, Related to Figure S3**

**Table S4. Crystallographic data collection and refinement statistics of Cas9 on-target and off-target complexes, Related to STAR methods**

**Table S5. Summary of non-canonical base pairing and nucleobase distortions observed in structural data, Related to Figure 1**

**Table S6. 3DNA 2.0 analysis of the helical parameters and sugar puckering of characterised on-target and off-target duplexes, Related to Figure 1**

**Table S7. Effect of Cas9 R895A mutation on off-target cleavage kinetics, Related to Figure S11**

Kinetic and thermodynamic parameters of *FANCF* on- and off-target substrate DNA cleavage by wild-type and R895A Cas9. Cleavage rate constants (k_obs_) were derived from single-exponential function fitting. Substrate binding and dissociation rate constants (k_on_ and k_off_) and the equilibrium dissociation constant (K_d_) were determined using a DNA nanolever (switchSENSE) binding assay. Intervals indicate standard error of mean.

**Table S8. Cleavage kinetics of off-targets with predicted PAM-distal deletions, Related to Figures S18 and S19**

Kinetic and thermodynamic parameters determined for *AAVS1* and *CD34* off-targets with predicted PAM-distal deletions. The apparent cleavage rate constants (k_obs_) were derived from a single-exponential function fitting of measured cleavage. Substrate binding (k_on_) and dissociation (k_off_) constants were determined using SwitchSENSE assay. Intervals indicate standard error of mean. NM indicates that values were not measured.

**Table S9. List of oligonucleotides used in this study, Related to STAR methods**

## Resource availability

### Lead contact

Further information and requests for resources and reagents should be directed to and will be fulfilled by the lead contact, Prof. Martin Jinek

### Material availability

This study did not generate new unique reagents.

### Data and code availability

- Coordinates have been deposited to the Protein Data Bank under the PDB accession numbers: 7QQO, 7QR7, 7QQP, 7QQQ, 7QQR, 7QQS, 7QQT, 7QQU, 7QQV, 7QQW, 7QQX, 7QR5, 7QQZ, 7QR8, 7QR0, 7QR1 and 7ZO1. Raw sequencing files for the SITE-Seq assay have been deposited to BioProject.
- This paper does not report original code.
- Any additional information required to reanalyze the data reported in this paper is available from the lead contact upon request.

### Experimental model and subject details

The *E. coli* strains Rosetta 2 (DE3) was used for recombinant protein expression for structural studies and biochemical *in vitro* experiments. T cells used for cellular editing experiments were isolated from human primary blood mononuclear cells were purchased from STEMCELL Technologies.

## METHODS

### DNA oligonucleotides and substrates

Sequences of DNA oligonucleotides used in this study are summarised in **Table S9**. Crystallisation substrates were synthesised by Sigma Aldrich without further purification, sgRNA transcription templates and ATTO-532-labelled cleavage substrates were synthesised by Integrated DNA Technologies, Inc., with PAGE and HPLC purification, respectively. Partially double stranded crystallisation substrates were prepared by mixing complementary oligonucleotides in a 1:1 molar ratio (as determined by 260 nm absorption), heating to 95 °C for 5 minutes and slow cooling to room-temperature. Cleavage substrates were prepared similarly, except that a 2-fold molar excess of the non-target strand was used.

### Cas9 protein expression and purification

*Streptococcus pyogenes* Cas9 wild type protein and the nuclease dead mutant (D10A, H840A) were both recombinantly expressed for 16 hours at 18 °C in *Escherichia coli* Rosetta 2 (DE3) (Novagen) N-terminally fused to a hexahistidine affinity tag, the maltose binding protein (MBP) polypeptide, and the tobacco etch virus (TEV) protease cleavage site. Cells were resuspended and lysed in 20 mM HEPES-KOH pH 7.5, 500 mM KCl, 5 mM imidazole, and supplemented with added protease inhibitors. Clarified lysate was loaded on a 10 ml Ni-NTA Superflow column (QIAGEN), washed with 7 column volumes of 20 mM HEPES-KOH pH 7.5, 500 mM KCl, 5 mM imidazole, and eluted with 10 column volumes of 20 mM HEPES-KOH pH 7.5, 250 mM KCl, 200 mM imidazole. Salt concentration is adjusted and protein is loaded on a 10 ml HiTrap Heparin HP column (GE Healthcare) equilibrated in 20 mM HEPES-KOH pH 7.5, 250 mM KCl, 1 mM DTT. The column is washed with 5 column volumes of 20 mM HEPES-KOH pH 7.5, 250 mM KCl, 1 mM DTT, and Cas9 is eluted with 17 column volumes of 20 mM HEPES-KOH pH 7.5, 1.5 M KCl, 1 mM DTT, in a 0-32% gradient (peak elution around 500 mM KCl). His_6_-MBP tag was removed by TEV protease cleavage overnight with gentle shaking. The untagged Cas9 was concentrated and applied to a Superdex 200 16/600 (GE Healthcare) and eluted with 20 mM HEPES-KOH pH 7.5, 500 mM KCl, 1 mM DTT. Purified protein was concentrated to 10 mg/ml, flash frozen in liquid nitrogen and store at −80 °C. DTT was omitted in the size-exclusion step of the purification when protein was used for switchSENSE measurements.

### sgRNA transcription and purification

sgRNAs are transcribed from a double stranded PCR product template amplified from a plasmid in a 5 ml transcription reaction (30 mM Tris-HCl pH 8.1, 25 mM MgCl_2_, 2 mM spermidine, 0.01% Triton X-100, 5 mM CTP, 5 mM ATP, 5 mM GTP, 5 mM UTP, 10 mM DTT, 1 µM DNA transcription template, 0.5 units inorganic pyrophosphatase (Thermo Fischer), 250 µg homemade T7 RNA polymerase. The reaction is incubated at 37 °C for 5 hours, and then treated for 30 minutes with 15 units of RQ1 DNAse (Promega). The transcribed sgRNAs are subsequently PAGE purified on an 8% denaturing (7 M urea) polyacrylamide gel, and lastly ethanol precipitated and resuspended in DEPC treated water.

### Crystallisation of Cas9 ternary complexes and structure determination

To assemble the Cas9 on-/off-target ternary complexes, the Cas9 protein is first mixed with the sgRNA in a 1:1.5 molar ratio and incubated at room temperature for 10 minutes. Next, the binary complex is diluted to 2 mg/ml with 20 mM HEPES-KOH 7.5, 250 mM KCl, 1 mM DTT, 2 mM MgCl_2_ buffer, pre-annealed 100 µM DNA substrate is added in a 1:1.8 molar ratio and the complex is incubated another 10 minutes at room temperature. For crystallisation, 1 µl of the ternary complex (1-2 mg/ml) is mixed with 1 µl of the reservoir solution (0.1 M Tris-acetate pH 8.5, 0.3-0.5 M KSCN, 17-19% PEG3350) and crystals are grown at 20 °C using the hanging drop vapour diffusion method. In some cases, microseeding was be used to improve crystal morphology. Crystals are typically harvested after 2-3 weeks, cryoprotected in 0.1 M Tris-acetate pH 8.5, 0.4 M KSCN, 30% PEG3350, 15% ethylene glycol, 1 mM MgCl_2_, and flash-cooled in liquid nitrogen. Diffraction data was obtained at beamlines PXI and PXIII of the Swiss Light Source (Paul Scherrer Institute, Villigen, Switzerland) and were processed using the XDS package (Kabsch, 2010). Structures were solved by molecular replacement through the Phaser module of the Phenix package (Adams et al., 2010) using the PDB ID: 5FQ5 model omitting the RNA-DNA target duplex from the search. Model adjustment and duplex building was completed using COOT software (Emsley et al., 2010). Atomic model refinement was performed using Phenix.refine (Adams et al., 2010). Protein-nucleic acid interactions were analysed using the PISA web server (Krissinel and Henrick, 2007). Characterisation of the guide-protospacer duplex was performed using the 3DNA 2.0 web server (Li et al., 2019). Structural figures were generated using PyMOL and ChimeraX (Pettersen et al., 2021).

### *In vitro* nuclease activity assays

Cleavage reactions were performed at 37 °C in reaction buffer, containing 20 mM HEPES pH 7.5, 250 mM KCl, 5 mM MgCl_2_ and 1 mM DTT. First, Cas9 protein was pre-incubated with sgRNA in 1:1.25 ratio for 10 minutes at room temperature. The protein-RNA complex was rapidly mixed with the target strand-ATTO-532-labelled dsDNA, to yield final concentrations of 1.67 μM protein and 66.67 nM substrate in a 7.5 µl reaction. Time points were harvested at 1, 2.5, 5, 15, 45, 90, 150 minutes, and 24 hours. Cleavage was stopped by addition of 2 µl of 250 mM EDTA, 0.5% SDS and 20 μg of Proteinase K. Formamide was added to the reactions with final concentration of 50%, samples were incubated at 95 °C for 10 minutes, and resolved on a 15% denaturing PAGE gel containing 7M urea and imaged using a Typhoon FLA 9500 gel imager. Depicted error bars correspond to the standard deviation from four independent cleavage reactions. Rate constants (k_obs_) were extracted from single exponential fits: [Product] = A*(1-exp(-k_obs_*t))

### switchSENSE analysis

The target strands (TS) containing a 3’ flanking sequence complementary to the ssDNA covalently bound to the chip electrode, and the non-target strands (NTS) (**Table S9**) were resuspended in a buffer containing 10 mM Tris-HCl pH 7.4, 40 mM NaCl, and 0.05% Tween 20. The matching TS:NTS duplex is pre-annealed and hybridised to the chip anode. The Cas9 protein was mixed with the sgRNAs at a 1:2 protein:RNA molar ratio, and the complex was incubated for 30 min at 37 °C in association buffer containing 20 mM HEPES-KOH pH 7.5, 150 mM KCl, 2 mM MgCl_2_, 0.01% Tween 20. All switchSENSE experiments were performed on a DRX analyser using CAS-48-1-R1-S chips (Dynamic Biosensors GmbH, Martinsried, Germany). Kinetics experiments were performed at 25 °C in association buffer, with an association time of 5 min, dissociation time of 20 min, and a flow rate of 50 µl/min.

### SITE-Seq assay

SITE-Seq assay reaction conditions were performed as described previously (Cameron et al., 2017). Briefly, high molecular weight genomic DNA (gDNA) was purified from human primary T cells using the Blood & Cell Culture DNA Maxi Kit (Qiagen) according to the manufacturer’s instructions. RNPs comprising the guides were biochemically assembled for gDNA digestion. Specifically, equal molar amounts of crRNA and tracrRNA were mixed and heated to 95 °C for 2 min then allowed to cool at room temperature for ~5 min. Three-fold molar excess of the guides were incubated with *Streptococcus pyogenes* Cas9 (SpCas9) in cleavage reaction buffer (20 mM HEPES pH 7.4, 150 mM KCl, 10 mM MgCl2, 5% glycerol) at 37 °C for 10 min. In a 96-well plate format, 10 µg of gDNA was treated with 0.2 pmol (4 nM), 0.8 pmol (16 nM), 3.2 pmol (64 nM), and 12.8 pmol (256 nM) of each RNP in 50 µL total volume in cleavage reaction buffer. Each cleavage reaction was performed in triplicate. Negative control reactions were assembled in parallel and did not include RNP. gDNA was treated with RNPs for 4 hours at 37 °C. SITE-Seq assay library preparation and sequencing was performed as described previously and the final library was loaded onto the Illumina NextSeq platform (Illumina, San Diego, CA), and ~1-3 M reads were obtained for each sample.

### SITE-Seq assay analysis and selection for cellular validation

SITE-Seq assay recovered off-targets were filtered for sites that had read-pileups proximal to the expected cut site, a PAM comprising at least one guanine base, fewer than 12 mismatches (reasoning that sites with 12 or more mismatches are likely spurious peaks not resulting from Cas9-induced double-strand breaks), and all sites with 11 mismatches were visually inspected and included in analysis if a putative deletion or insertion would result in a reduction of at least 4 mismatches relative to the spacer sequence.

### *In silico* mismatch, deletion, and insertion classification

Predictive classification of SITE-Seq assay recovered off-target sites as pure mismatches, deletions, or insertions was executed using a scoring algorithm which consisted of the following sequential steps (**Figure S15**):

i. For each off-target, a gap library was generated where a single nucleotide gap was introduced between each nucleotide in the off-target sequence.
ii. The off-target gap library was then aligned to the spacer sequence and each alignment was scored based on the number of matched bases between the spacer and gapped off-target pair. If the gapped off-target with the highest alignment score improved alignment by at least 4 nucleotides relative to the non-gapped spacer–off-target alignment, the off-target sequence was marked as a single-nucleotide deletion and removed from subsequent analysis.
iii. The remaining pool of off-targets were then aligned to a spacer gapped library where a single nucleotide gap was introduced at each positing in the spacer.
iv. The spacer gap library was then aligned to each off-target sequence and each alignment was scored based on the number of matched bases between the off-target and the gapped spacer pair. If the gapped spacer with the highest alignment score improved alignment by at least 4 nucleotides relative to the non-gapped spacer–off-target alignment, the off-target sequence was marked as a single-nucleotide insertion and removed from subsequent analysis.
v. The remaining off-target for which the spacer–off-target alignment was not improved by single-nucleotide deletions or insertions were annotated as a mismatched off-target.

### Generation of ribonucleoprotein complexes (RNP) for electroporation

Synthetic crRNA and tracrRNA oligonucleotides were ordered from Integrated DNA Technologies (Coralville, IA) as AltR^TM^ reagents. Ribonucleoprotein complexes (RNPs) were formulated by incubating 480 pmols of each crRNA and tracrRNA and heated to 95 °C for 2 min then allowed to cool at room temperature for ~5 min. Annealed guides were then mixed with 160 pmols of Cas9 protein (1:3 molar ratio of Cas9 to guides) in RNP assembly buffer (20 mM HEPES pH 7.4, 150 mM KCl, 10 mM MgCl_2_, 5% glycerol) and incubated at 37 °C for 10 min. RNP were then serial diluted two-fold by mixing with equal volume of RNP assembly buffer across seven concentrations down to 2.5 pmol of Cas9 and 7.5 pmols of crRNA-tracrRNA. RNPs were kept at 4 °C or on ice until transfection.

### T cell handling and transfections

T cells were isolated from peripheral blood mononuclear cells (PBMCs) using the RoboSep-S cell isolation platform (STEMCELL Technologies) with EasySep Human T cell Isolation Kit (STEMCELL Technologies). Cells were then activated for 3 days in “Complete media” (Immunocult-XF T cell expansion media (STEMCELL Technologies), CTS Immune Cell serum replacement (Fisher Scientific), 1x antibiotic-antimycotic (Fisher Scientific), and recombinant human Interleukin-2 (rhIL-2, 100 units/mL)) in the presence of anti-CD3/CD28 Dynabeads (Fisher Scientific) at a bead:cell ratio of 1:1. On the third day, beads were removed and cells expanded in Complete media for 24 hours. Cells were then harvested, washed once with phosphate-buffered saline (PBS) and resuspended at 10^7^ cells/mL in P4 Primary Cell Nucleofector Solution (Lonza). 20 µL of resuspended cells (corresponding to ~ 200,000 cells) were then combined with 2.5 µL of RNP and electroporated using the Lonza 4D 96-well Nucleofector electroporation system using pulse code CA137. Transfected cells were then recovered in 180 µL/sample of Complete media. After 48 hours, the electroporated T cells were pelleted and gDNA was isolated by adding 50 μL/well QuickExtract DNA extraction solution (Epicentre), followed by incubation at 37 °C for 10 minutes, 65 °C for 6 minutes, and 95 °C for 3 minutes. The isolated gDNA was diluted with 50 μL sterile water to achieve ~ 2,000 genome equivalents/μL and samples were stored at −20 °C until NGS analysis.

### Targeted amplicon sequencing

Cas9 on- and off-target sites were amplified in a two-step PCR reaction. In brief, 3.75 μl (corresponding to ~7,500 cells) of lysate was used as a template for PCR amplification with Q5 Hot-Start High Fidelity DNA Polymerase (NEB) and unique primer pairs containing an internal locus-specific region and an outer Illumina-compatible adapter sequence. A second PCR reaction targeting the outer-adapter sequence was performed to append unique indices to each amplicon. Sites were sequenced on a MiSeq with 2 × 151 paired-end reads and v2 chemistry, or NextSeq with 2 × 151 paired-end reads and v2 or v2.5 chemistry (Illumina), and aligned to the hg38 genomic assembly. For each site, indels were tallied if they occurred within 3 nucleotides of the putative Cas9 cut site, and editing efficiencies were calculated by subtracting the percentage of indels in untransfected cells from the percentage of indels in RNP-transfected cells. Depth of coverage was ~5,000–20,000 reads per amplicon, and all samples with <500 reads aligning to the predicted amplicon were discarded. Lower limit of detection (*e.g.,* ≥0.1%) was determined by titration of NIST genomic standards and assessment of expected versus measured values (Data not shown).

### Molecular Dynamics (MD) simulations

MD simulations were performed using a well-established protocol, previously employed in several computational studies of CRISPR-Cas9 (Mitchell et al., 2020; Ricci et al., 2019). The Amber ff14SB force field (Maier et al., 2015) was adopted, including the ff99bsc1 corrections for DNA (Ivani et al., 2016) and the ff99bsc0+χOL3 corrections for RNA (Banas et al., 2010; Zgarbova et al., 2011). Explicit water molecules were described using the TIP3P model (Jorgensen et al., 1983), while the Li & Merz model was used for Mg2+ ions (Li and Merz, 2014). An integration time step of 2 fs was applied. All bond lengths involving hydrogen atoms were constrained using the SHAKE algorithm. Temperature control (300 K) was performed via Langevin dynamics (Turq et al., 1977), with a collision frequency γ = 1. Pressure control was accomplished by coupling the system to a Berendsen barostat (Berendsen et al., 1984), at a reference pressure of 1 atm and with a relaxation time of 2 ps. The systems were subjected to energy minimization to relax water molecules and counter ions, keeping the protein, the RNA, DNA and Mg2+ ions fixed with harmonic position restraints of 300 kcal/mol · Å2. Then, the systems were heated up from 0 to 100 K in the canonical ensemble (NVT), by running two simulations of 5 ps each, imposing position restraints of 100 kcal/mol · Å2 on the above-mentioned elements of the system. The temperature was further increased up to 200 K in ~100 ps of MD in the isothermal-isobaric ensemble (NPT), reducing the restraint to 25 kcal/mol · Å2. Subsequently, all restraints were released, and the temperature of the systems was raised up to 300 K in a single NPT simulation of 500 ps. After ~ 1.1 ns of equilibration, ~10 ns of NPT runs were carried out allowing the density of the systems to stabilize around 1.01 g cm-3. Finally, production runs were carried out in the NVT ensemble, collecting ~500 ns in three replicates for each of the systems. These simulations have been performed using the GPU-empowered version of AMBER 20 (D.A. Case, 2021; Salomon-Ferrer et al., 2013). Analysis of the RNA:DNA conformational dynamics has been done using the CURVES+ code (Lavery et al., 2009). Molecular simulations were based on three X-ray structures of CRISPR-Cas9: TRAC on-target (PDB: 7OX8), AAVS1 on-target #2 and FANCF on-target, displaying 20 base pair long RNA:TS DNA hybrid and a cleaved NTS. To study the accommodation of rA-dA mismatches at various positions along the heteroduplex, we systematically mutated the base pairs to rA-dA mismatches (TRAC on-target for positions 1, 11, 15, and 20; FANCF on-target for positions 3-5, 9, and 18; AAVS1 on-target for positions 2, 6-8, 10, 12-14, 16, 17, and 19), using the LEaP tool in the AMBER 20 code (Lavery et al., 2009). Additionally, the pre-cleavage state crystal structure of CRISPR-Cas9 with no mismatches (PDB: 4UN3) was also simulated for comparison. The obtained model systems were embedded in explicit waters, and counterions were added to neutralize the total charge at physiological conditions, leading to periodic simulation cells of ~145*115*150 Å3 and a total of ~220,000 atoms for each system.

